# The genetic basis of structural colour variation in mimetic *Heliconius* butterflies

**DOI:** 10.1101/2021.04.21.440746

**Authors:** Melanie N. Brien, Juan Enciso Romero, Victoria J. Lloyd, Emma V. Curran, Andrew J. Parnell, Carlos Morochz, Patricio A. Salazar, Pasi Rastas, Thomas Zinn, Nicola J. Nadeau

**Affiliations:** Ecology and Evolutionary Biology, School of Biosciences, The University of Sheffield, Alfred Denny Building, Western Bank, Sheffield, S10 2TN, United Kingdom; Biology Program, Faculty of Natural Sciences, Universidad del Rosario, Bogotá, Colombia; Department of Physics and Astronomy, The University of Sheffield, Hicks Building, Hounsfield Road, Sheffield, S3 7RH, United Kingdom; Biology & Research Department, Mashpi Lodge, Ecuador; Institute of Biotechnology, University of Helsinki, Finland; ESRF - The European Synchrotron, 38043, Grenoble Cedex 9, France

**Keywords:** Structural colour, iridescence, gene expression, *Heliconius*, QTL, convergence

## Abstract

Structural colours, produced by the reflection of light from ultrastructures, have evolved multiple times in butterflies. Unlike pigmentary colours and patterns, little is known about the genetic basis of these colours. Reflective structures on wing-scale ridges are responsible for iridescent structural colour in many butterflies, including the Müllerian mimics *Heliconius erato* and *Heliconius melpomene*. Here we quantify aspects of scale ultrastructure variation and colour in crosses between iridescent and non-iridescent subspecies of both of these species and perform quantitative trait locus (QTL) mapping. We show that iridescent structural colour has a complex genetic basis in both species, with offspring from crosses having wide variation in blue colour (both hue and brightness) and scale structure measurements. We detect two different genomic regions in each species that explain modest amounts of this variation, with a sex-linked QTL in *H. erato* but not *H. melpomene*. We also find differences between species in the relationships between structure and colour, overall suggesting that these species have followed different evolutionary trajectories in their evolution of structural colour. We then identify genes within the QTL intervals that are differentially expressed between subspecies and/or wing regions, revealing likely candidates for genes controlling structural colour formation.

## En español

El color estructural, producto del reflejo de la luz en ultraestructuras, ha evolucionado múltiples veces en mariposas. A diferencia de los patrones de color pigmentario, se sabe poco sobre la genética del color estructural. Las estructuras reflectivas en las crestas de las escamas de las mariposas son responsables del color estructural iridiscente en varias especies de mariposas, incluyendo a las co-miméticas *H. erato* y *H. melpomene*. En este artículo cuantificamos aspectos de la variación de la ultraestructura y el color estructural en cruces entre sub-especies iridiscentes y melánicas de ambas especies y hacemos mapeo de loci de rasgos cuantitativos (QTL). Mostramos que el color estructural iridiscente tiene una base genética compleja en ambas especies; la progenie de ambos cruces varía ampliamente en color (tono y brillo) y en la morfología de sus escamas. Detectamos dos regiones genómicas diferentes en cada especie que explican cantidades modestas de esta variación, con un QTL ligado al sexo en *H. erato* pero no en *H. melpomene*. También encontramos diferencias entre especies en las relaciones entre estructura y color, lo que en general sugiere que estas especies han seguido trayectorias diferentes en la evolución del color estructural. Identificamos genes al interior de los intervalos de confianza de los QTL que se expresan diferencialmente entre sub-especies y regiones del ala, revelando posibles genes candidatos que controlan la formación del color estructural.

## Introduction

Structural colours are some of the most vivid and striking colours found in nature. They are formed from the reflection and refraction of light from physical ultrastructures and examples of these can be found in nearly all groups of organisms. The structural colours of butterflies and moths are among the best described and play diverse roles, including initiation of courtship and mating behaviour (Silberglied and Taylor, 1978; Obara *et al*., 2008), sex and species discrimination (Rutowski, 1977), long distance mate recognition (Sweeney, Jiggins and Johnsen, 2003) and signalling of quality and adult condition (Kemp and Rutowski, 2007).

Butterflies and moths have evolved several mechanisms of structural colour production by modifying different components of wing scale morphology (Ghiradella, 1991). Scales typically consist of a flat lower lamina connected to an upper lamina by pillar-like trabeculae, with a small space separating the upper and lower laminae (Figure 1). The lower lamina can act as a thin film reflector that produces hues ranging from violet to green depending on its thickness (Stavenga, Leertouwer and Wilts, 2014; Wasik *et al*., 2014; Thayer, Allen and Patel, 2020). The upper lamina has a more complex structure; it consists of a parallel array of ridges connected by cross-ribs, and modifications to these can yield diverse optical effects. For example, a lamellar structure in the ridges forms multilayer reflectors that produce the iridescent (angle dependent) blue in *Morpho* butterflies (Vukusic *et al*., 1999) and UV reflectance in *Colias eurytheme* (Eisner *et al*., 1969; Ghiradella, 1974). The variations in hue and brightness of colour produced in the intricate structures of the upper lamina depend on an interplay between the number of lamellae, the thickness of each layer and the spacing between the ridges (Parnell *et al*., 2018).

**Figure 1.**
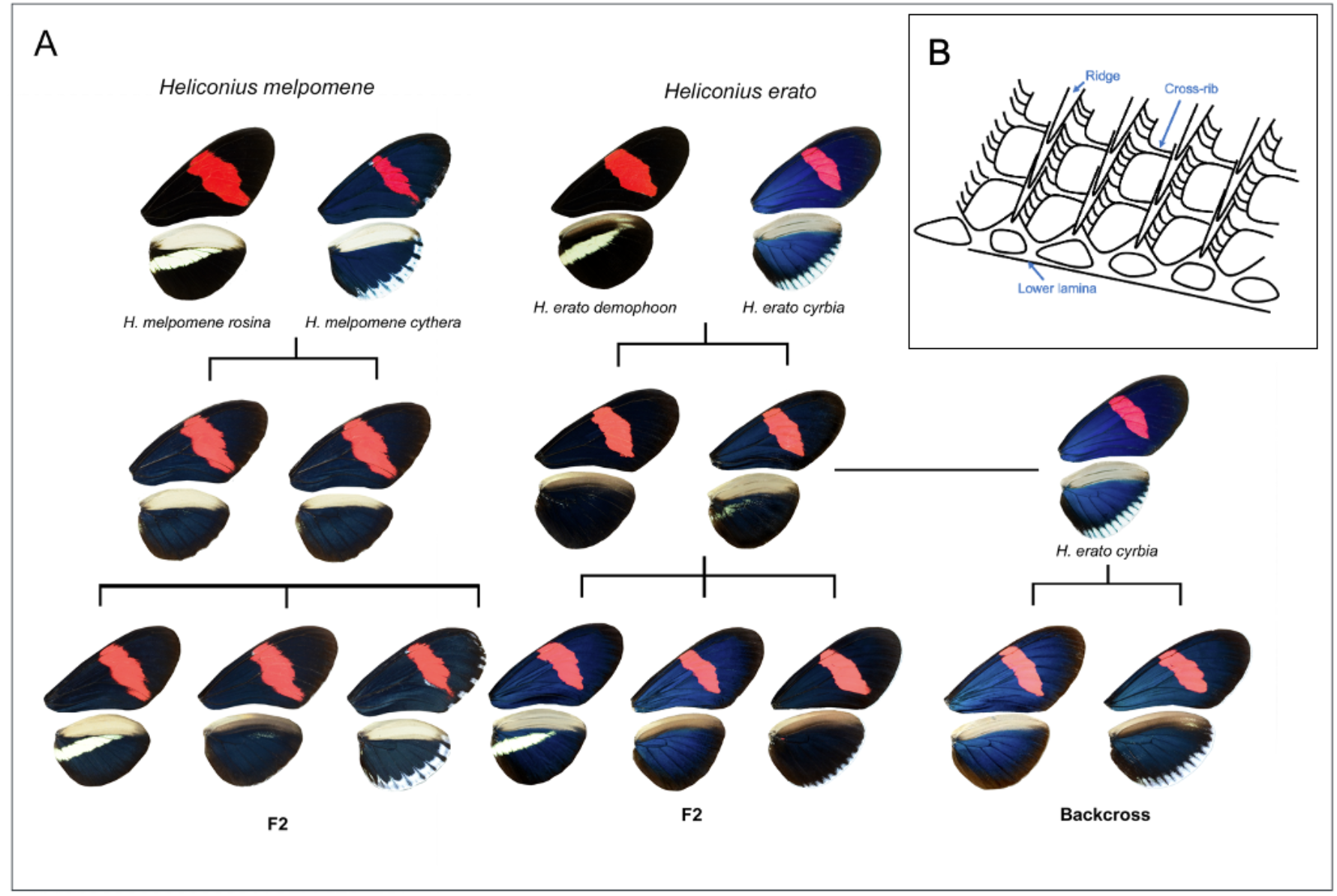
(A) Crosses between iridescent and non-iridescent morphs of *Heliconius melpomene* and *Heliconius erato*. For *H. melpomene* we used F2 crosses, plus one cross thought to be F1xF2 (not shown). For *H. erato*, we used F2 crosses and a backcross to the iridescent subspecies. (B) Scales are formed of a lower lamina and an upper lamina, which is made up of longitudinal ridges connected by cross-ribs.

Recent studies have begun to uncover the genetic and developmental basis of structural colours in some species (Lloyd and Nadeau, 2021), revealing a common pattern in *Bicyclus anynana* and *Junonia coenia;* artificial selection for colourful phenotypes quickly resulted in changes in lower lamina thickness, and consequently hue, in a relatively small number of generations (Wasik *et al*., 2014; Thayer, Allen and Patel, 2020). Knockouts of known colour pattern genes (Concha *et al*., 2019), and genes involved in pigment synthesis pathways (Zhang, Mazo-Vargas and Reed, 2017; Matsuoka and Monteiro, 2018), have shown that modification of these can result in altered scale ultrastructure, and moreover has brought about unexpected instances of structural colour (Zhang, Mazo-Vargas and Reed, 2017). Interestingly, there are groups of butterflies for which the gene *optix*, a known major colour pattern gene (Reed *et al*., 2011), can jointly control pigment based colouration and thickness of the lower lamina, producing blue structural colour (Thayer, Allen and Patel, 2020). It remains unclear whether this joint control extends to other butterfly taxa that produce structural colour using lower lamina reflection. In species with ridge reflectors, such as *Heliconius*, this does not seem to be the case (Zhang, Mazo-Vargas and Reed, 2017). More efforts are required to understand the genetic underpinnings and evolution of upper lamina architecture and the organisation of reflective nanostructures in the scales of iridescent butterflies.

Wing colour patterns have been widely studied in the *Heliconius* butterflies, a group of butterflies with a diverse set of aposematic colour patterns. These patterns show examples of both convergent evolution between distantly related species, and divergent evolution within species. Some species form mimicry rings, in which wing patterning is under strong positive frequency-dependent selection due to predation (Mallet and Barton, 1989). Pigment colour patterns are largely determined by a small number of genes which are homologous across species. Extensive research has uncovered a toolkit of five loci which control much of the colour pattern variation in *Heliconius* species, and some other Lepidoptera (Nadeau, 2016). *Heliconius* also display structural colour, and in comparison to the well-studied pigmentary colours, very little is known about the development and genetic basis of these. While overall scale morphology is similar between iridescent and non-iridescent scales in *Heliconius*, those with blue structural colour have overlapping ridge lamellae which act as multilayer reflectors (as in *Morpho*), along with a greater density of ridges on the scale (narrower ridge spacing) (Brien *et al*., 2018; Parnell *et al*., 2018).

Structural colour has evolved multiple times within the *Heliconius* genus (Parnell *et al*., 2018). In some species, all subspecies have iridescent colour, while others exhibit interspecific variation in iridescence. *Heliconius erato* and *Heliconius melpomene* are two co-mimicking species which diverged around 10-13 Mya (Kozak *et al*., 2015) with each evolving around 25 different colour pattern morphs (Sheppard *et al*., 1985). Most of the different colour patterns are produced by pigment colours, but subspecies found west of the Andes in Ecuador and Colombia also have an iridescent blue structural colour. *H. erato cyrbia* and *H. melpomene cythera* found in Western Ecuador have the brightest iridescence, while subspecies *H. erato demophoon* and *H. melpomene rosina*, found to the north in Panama, are matt black in the homologous wing regions (Figure 1). A hybrid zone forms between the iridescent and non-iridescent groups where they meet near the border between Panama and Colombia, and here, populations with intermediate levels of iridescence can be found (Curran *et al*., 2020). Continuous variation in iridescent colour is observed in the centre of the hybrid zone and in experimental crosses (Brien *et al*., 2018), suggesting that this trait is controlled by multiple genes. The evolution of pigmentation and simple colour pattern traits have frequently been shown to involve the re-use of a small number of genes across animal species (Hubbard *et al*., 2010; Manceau *et al*., 2010; Nadeau, 2016). However, we may expect the genetic basis of a quantitative trait controlled by multiple genes, such as iridescence in these species, to be less predictable (Conte *et al*., 2012). In addition, iridescence in *H. e. cybria* is much brighter than in *H. m. cythera* (Parnell et al., 2018), suggesting some differences in scale structure and presumably genetic control of this structure formation process.

Here, we use crosses between subspecies of iridescent and non-iridescent *Heliconius* to determine the genetics of both colour and scale ultrastructure traits for the first time. We measure the intensity of blue colour and overall luminance (brightness) to assess variation in colour. We complement our estimates of colour variation with high throughput measurements of ridge spacing and cross-rib spacing using ultra small-angle X-ray scattering (USAXS). Using a quantitative trait locus (QTL) mapping approach, we can identify the location and effect sizes of loci in the genome that are controlling variation in iridescent colour. We then use RNA sequencing data from the same subspecies of each species to identify genes that are differentially expressed, both between subspecies and between wing regions that differ in scale type. Comparison of the genetic basis of these traits between *H. melpomene* and *H. erato*, two distantly related mimetic species, allows us to ask whether, like pigment colour patterns, variation in iridescent colour and scale structure is also an example of gene reuse.

## Methods

### Experimental crosses

Experimental crosses were performed using geographical morphs of both *Heliconius erato* and *Heliconius melpomene*. In both species, morphs from Panama (*H. e. demophoon* and *H. m. rosina*) were crossed with morphs from Western Ecuador (*H. e. cyrbia* and *H. m. cythera*), then the F1 generation crossed with each other to produce an F2. For *H. erato*, we also analysed a backcross between the F1 and *H. e. cyrbia* (Figure 1). Due to a mix-up in the insectary, one of our largest *H. melpomene* broods, named ‘EC70’, was obtained from a cross between an F1 father and a mother of unknown parentage, likely an F2 individual. Further details of the crosses are in Table S1. A total of 155 *H. erato* individuals from 5 broods were used to generate linkage maps and perform QTL mapping (3 *demophoon* and 3 *cyrbia* grandparents, 11 F1 parents, and 40 backcross and 99 F2 offspring). For *H. melpomene*, data from 4 broods made up of 228 individuals were used (1 *rosina* and 2 *cythera* grandparents, 6 parents and 219 offspring, Table S1). Some of these crosses have previously been used for an analysis of quantitative pattern variation (Bainbridge *et al*., 2020). Details of sequencing and linkage map construction are given in Bainbridge et al. (Bainbridge *et al*., 2020) and in the Supplementary Material.

### Phenotypic measurements

In the offspring of these crosses, we measured four phenotypes - blue colour (BR), luminance, ridge spacing and cross-rib spacing. Wings were photographed under standard lighting conditions (full details in (Brien *et al*., 2018)). A colour checker in each photograph was used to standardise the photographs using the levels tool in Adobe Photoshop (CS3). RGB values were extracted from two blue/black areas of each wing (proximal areas of both the forewing and hindwing, Figure S1) and averaged. Blue-red (BR) values were used as a measure of blue iridescent colour. These were calculated as (B-R)/(B+R), where 1 is completely blue and −1 is completely red. Luminance measured overall brightness and was calculated as R+G+B, with each colour having a maximum value of 255.

Scale structure measurements were extracted from ultra small-angle X-ray scattering (USAXS) data, from a single family of each species (n=56 *H. erato* F2 and n=73 *H. melpomene* (mother of unknown ancestry)). We measured between 33 and 113 points per individual along a linear proximodistal path across the proximal part of the forewing, which has the most vivid iridescence in the blue subspecies (Figure S1). The raw images were corrected for dark current and spatial distortion. SEM data from a subset of individuals was used to interpret the scattering patterns and develop robust methods for extracting mean ridge and cross-rib spacing values for the dorsal wing scales of all individuals (see Supplementary Material for details).

### Quantitative Trait Locus mapping

The R package R/qtl was used for the QTL analysis (Broman *et al*., 2003). For *H. erato*, initially the F2 crosses were analysed together and the backcross analysed separately. Genotype probabilities were calculated for these two groups using *calc.genoprob*. We ran standard interval mapping to estimate LOD (logarithm of the odds) scores using the *scanone* function with the Haley-Knott regression method. In the F2 analysis, sex and family were included as additive covariates, and family was included as an interactive covariate, to allow multiple families to be analysed together. Sex was included as a covariate in the backcross analysis to account for any sexual dimorphism. To determine the significance level for the QTL, we ran 1000 permutations, with *perm.Xsp=T* to get a separate threshold for the Z chromosome. A single F2 family (n=56) was used to analyse scale structure variation (ridge spacing and cross-rib spacing) using the same method, albeit a higher number of permutations was used for determining the significance level of the QTL (4000). For analyses of BR colour and luminance, LOD scores for the F2 crosses and the backcross were added together, to allow analysis of all individuals together to increase power, and the significance level recalculated in R/qtl.

Confidence intervals for the positions of QTL were determined with the *bayesint* function and we used a *fitqtl* model to calculate the phenotypic variance that each QTL explained. Genome scan plots and genotype plots were made with R/qtl2 (Broman *et al*., 2019). Genetic distances in the QTL results are based on the observed recombination rate and expressed in centimorgans (cM), which is the distance between two markers that recombine once per generation. These were related to physical distances based on the marker positions in the assembled reference genome of each species.

The same method was used to run genome scans for BR colour and luminance in *H. melpomene*. Since the parentage of the mother of the EC70 brood is unknown, the maternal alleles in the offspring could not be assigned as being from either a *cythera* or a *rosina* grandparent. Therefore, in this family only paternal alleles were taken into account (and all maternal alleles were assigned to a *rosina* grandparent), and the cross was treated as if a backcross. LOD scores of the three F2 families were added to the LOD score from the EC70 family, as in *H. erato*, and the significance level recalculated. Again, a single family was used for analysis of scale structures (EC70, n=73).

### Gene expression analysis

RNA sequence data was generated from 32 *H. erato* pupal wing samples (16 *H. e. demophoon*, 16 *H. e. cyrbia*) and *H. melpomene* pupal wing samples (16 *H. m. rosina*, 16 *H. m. cythera*), with individuals sampled from the same captive populations as those used for the crosses. Each of these samples contained 2 wing regions (the anterior hind-wing or “androconial” region, which has a different scale type, was dissected from the rest of the wing and sampled separately, Figure S1), and two developmental stages, 5 days post pupation (DPP) (50 % total pupation time) and 7 DPP (70 % total pupation time). Overall this gave four biological replicates for each tissue type/developmental stage/subspecies combination (Table S2).

Quality-trimmed reads were aligned to the respective *Heliconius* reference genomes using HISAT2 (version 2.1.0). Clustering of samples by Multi-Dimensional Scaling (MDS) on expression levels revealed one of the *H. m. rosina* individuals had been incorrectly labelled (which was also confirmed by analysis of nucleotide variants) and was removed from subsequent analyses. Each species was analysed separately to identify genes that were differentially expressed between subspecies and between the wing regions for the iridescent blue subspecies (Figure S1), using the quasi-likelihood (QL) F-test in R/Bioconductor package EdgeR (version 3.28.1). For the wing region comparison, we used a general linear model approach, with the two wing regions nested within “individual ID” for each individual. We then determined if any significantly differentially expressed genes (between subspecies or wing region) were within the mapped QTL intervals. We further determined if any genes were differentially expressed in parallel between species. Details of further analyses of these data including gene set enrichment analysis are given in the Supplementary Material.

## Results

### QTL mapping in H. erato

We found significant correlations between scale structure and colour measurements: ridge spacing is negatively correlated with both luminance and BR values (Figure S2). Cross-rib spacing is positively correlated with ridge spacing and also negatively correlates with BR values (supplementary text).

Significant QTL were found for three phenotypes in *H. erato* - BR colour, luminance and ridge spacing (Figure 2, Table 1, Table S3). When analysing the colour measurements, F2 and backcross genome scans were combined, and for BR values these showed 2 significant QTL on chromosomes 20 and the Z sex chromosome. These QTL were also found when analysing the F2 broods separately from the backcross brood (Figure S3). At both markers, individuals with Panama-type genotypes (Pan/Pan and Pan(W)) had lower BR values than Ecuador-type and heterozygous genotypes, following the expected trend (Figure 2). The QTL on the Z chromosome explained the largest proportion of the phenotypic variation in BR colour in both the F2 crosses (19.5%) and the backcross (24.6%), and the chromosome 20 QTL explained a further 12.3% in the F2 crosses.

**Figure 2.**
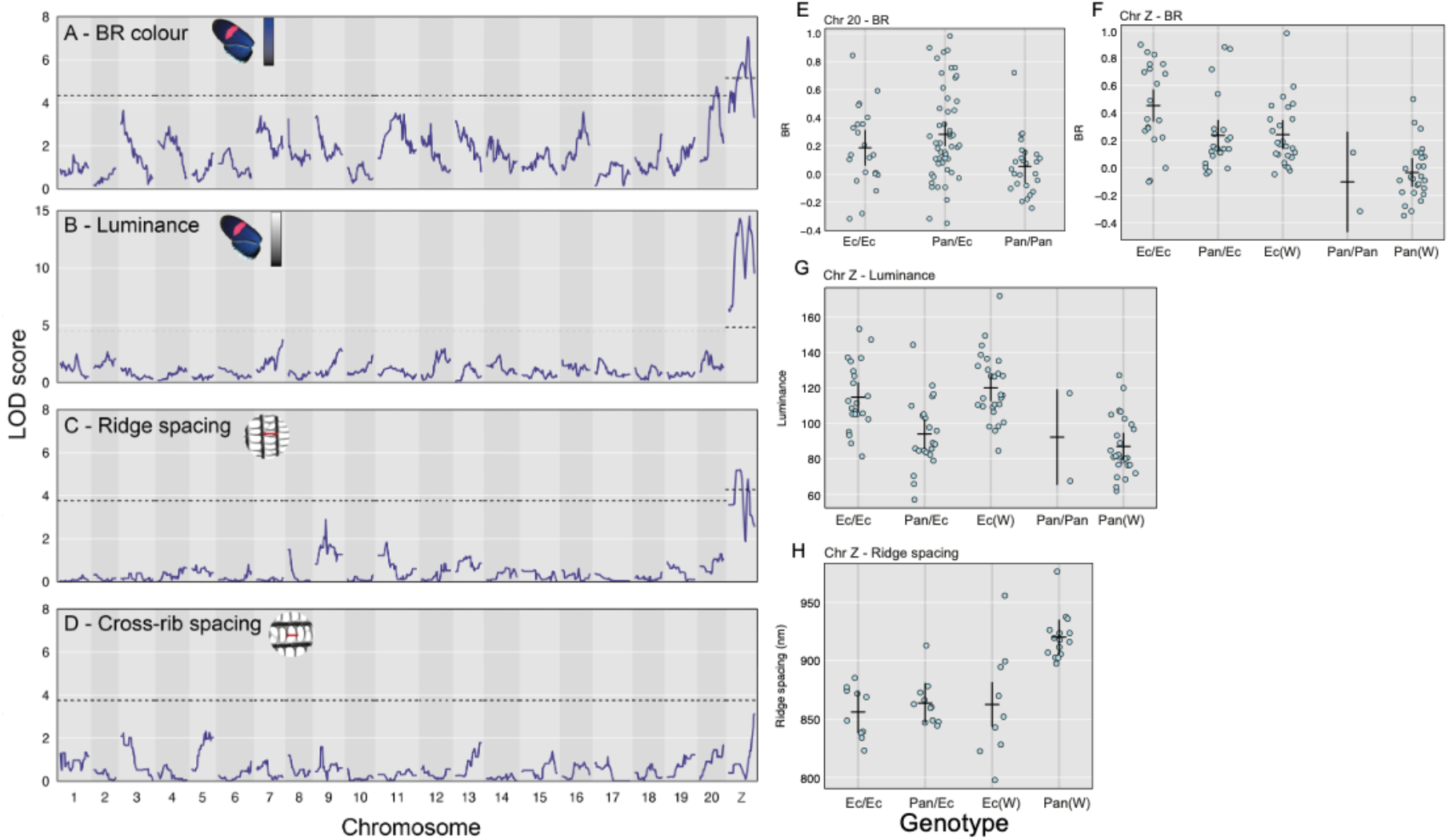
*H. erato* QTL plots including all families for BR colour (A) and luminance (B). QTL plots for a single family for ridge spacing (C) and cross-rib spacing (D). Dotted line shows p=0.05 significance level. The phenotypes of F2 individuals with different genotypes at the most significant markers within each *H. erato* QTL. The QTL for BR on the chromosome 20 (E) and the Z (F). Significant Z markers for luminance (G) and ridge spacing (H). ‘Pan’ denotes alleles from the Panama subspecies *demophoon*, and ‘Ec’ the Ecuador subspecies *cyrbia*. Only two individuals have homozygous Panama-type *demophoon* genotypes at the Z chromosome marker due to the small number of individuals with a *demophoon* maternal grandfather (Table S1). Marker positions are shown in Table 1.

**Table 1:** Significant QTL found for three phenotypes in *H. erato* and *H. melpomene*.

| Phenotype | Marker | Chromosome | Position (cM) | LOD | p |
| --- | --- | --- | --- | --- | --- |
| *Heliconius erato* |  |  |  |  |  |
| BR colour (all families) | Herato2101_12449252 | Z | 38.0 | 7.07 | 0.001 |
|  | Herato2001_12633065 | 20 | 32.9 | 4.75 | 0.022 |
| Luminance (all families) | Herato2101_12449398 | Z | 41.6 | 14.50 | <0.001 |
| Ridge Spacing (single family) | Herato2101_7491127 | Z | 23.0 | 5.21 | 0.013 |
| *Heliconius melpomene* |  |  |  |  |  |
| BR (all families) | Hmel203003o_2119654 | 3 | 15.22 | 7.26 | 0.001 |
| Luminance (all families) | Hmel203003o_2635435 | 3 | 17.97 | 13.61 | <0.001 |
| Ridge spacing (EC70) | Hmel207001o_11550301 | 7 | 53.61 | 5.71 | <0.001 |

Luminance (overall brightness of the wing region) was highly associated with the Z chromosome (Figure 2B). The significant marker did not map exactly to the same position as for the BR values but was apart by only 3.6cM, and confidence intervals for each overlap. Individuals with Ecuador-type alleles had higher luminance values than those with Panama-type alleles, showing the same trend as the BR values (Figure 2G). This QTL explained 40.2% of the variance in luminance values in the F2 crosses and 24.2% in the backcross. This was the only significant QTL for luminance, with nothing appearing on chromosome 20.

A single QTL on the Z chromosome was also significant for ridge spacing (Figure 2C). This marker was at a different position to the markers for BR and luminance, but mapped to the same marker as luminance when using the same individuals (Figure S3). All genotypes with one or two Ecuador-type alleles had similar ridge spacing, but those with a hemizygous Panama-type genotype (‘Pan(W)’ in Figure 2H) had significantly wider ridge spacing. This QTL explained 34.8% of variance in ridge spacing in this family. No significant QTL were found for cross-rib spacing, although the highest LOD score was seen on the Z chromosome (Figure 2D).

### QTL mapping in H. melpomene

In contrast to *H. erato*, scale structure measurements in *H. melpomene* did not correlate with either of the colour measurements (Figure S2, supplementary text). A single significant QTL for BR colour was found on chromosome 3 (Figure 3A, Table 1, Table S4) when combining the F2 families with EC70 (and for EC70 only, Figure S4). The marker explains 15.3% of phenotypic variation in EC70 (which should be an underestimate due to all maternal alleles being ignored) and 9.2% in the three F2 families. Luminance was also strongly associated with markers on chromosome 3 (Figure 3B, Figure S4). The associated marker was 2.75cM from the marker for BR colour, and the confidence intervals overlap. In contrast, for ridge spacing we found a significant QTL on chromosome 7 (using just the EC70 brood), explaining 30.3% of variation (Figure 3G). Again, no significant QTL were found for cross-rib spacing (Figure 3D). These results were generally supported by a genome-wide association analysis using all SNP variation (which allowed maternal variation in EC70 to be included) and did not reveal any additional loci (Figure S5, see Supplementary Material for full results and methods).

**Figure 3.**
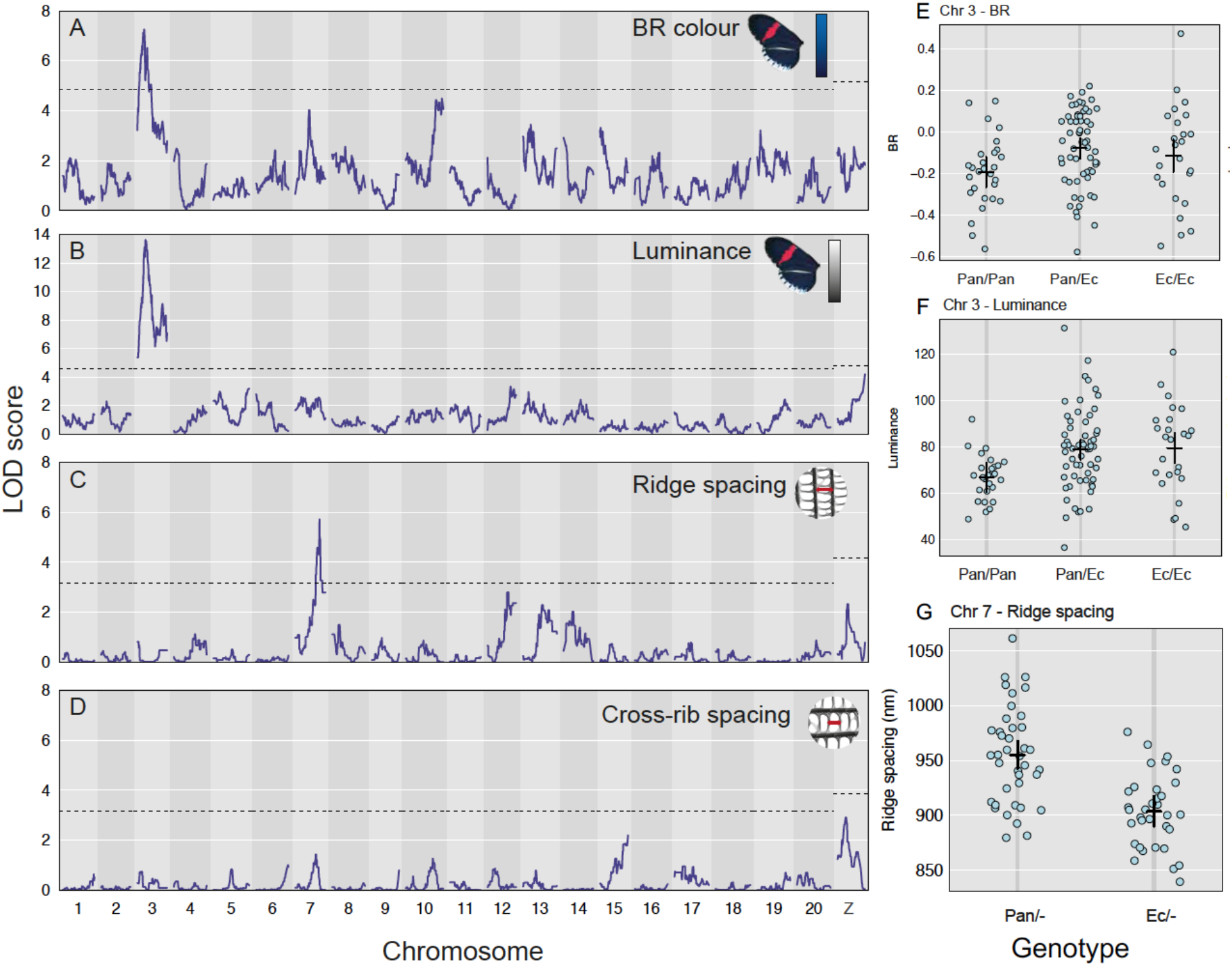
*H. melpomene* QTL plots including all families for BR colour (A) and luminance (B). QTL plots for a single family for ridge spacing (C) and cross-rib spacing (D). Genotype x phenotype plots for significant *H. melpomene* QTL. For the F2 families: (E) BR colour and (F) luminance. ‘Pan’ denotes alleles from the Panama subspecies *rosina*, and ‘Ec’ the Ecuador subspecies *cythera*. (G) Ridge spacing in the EC70 brood. Here we only know the origin of the paternal allele. Marker positions shown in Table 1.

Individuals with homozygous Panama-type genotypes at the mapped chromosome 3 markers had lower BR and luminance values (Figure 3). Individuals carrying Ecuador-type alleles at the mapped chromosome 7 marker showed reduced ridge spacing, consistent with the observation that the Panama subspecies has greater ridge spacing.

### Differential expression

A total of 24,118 genes were expressed in the wings of *H. erato* and 30,721 in the wings of *H. melpomene*. In both *H. erato* and *H. melpomene*, multidimensional scaling (MDS) analysis of expression levels revealed strong clustering by stage (dimension 1) and subspecies (dimension 2), leading to four distinct clusters (Figure S6). 907 and 1043 genes were differentially expressed (DE, FDR<0.05) between *H. erato cyrbia* and *H. erato demophoon* at 5 and 7 days post pupation, respectively (Table S5, S6). In *H. melpomene*, 203 and 29 genes were DE between *H. m. cythera* and *H. m. rosina* at 5 and 7 DPP, respectively (Table S7, S8). Comparing between wing regions, in *H. erato cyrbia* there was one gene at 5 DPP and 70 genes at 7 DPP DE (Table S9, S10); in *H. melpomene cythera*, there were six genes at 5 DPP and 50 genes at 7 DPP DE (Table S11, S12). Much of this DE will be due to the genome-wide divergence between subspecies (which is greater in *H. erato* than in *H. melpomene*, (Parnell *et al*., 2018; Curran *et al*., 2020)), we therefore used further comparisons to narrow down these lists of genes.

We may expect that genes involved in scale structure regulation would be DE both between subspecies and wing regions that differ in scale structure, but very few genes were found in both sets (Figure S7, Table S13). At 7 DPP, there were two genes upregulated in *H. erato* in both comparisons, *chitin deacetylase 1*, with a likely function in the deacetylation of chitin to chitosan and with potential structural roles in the cuticle (Thurmond *et al*., 2018). The other gene had similarity with the circadian clock-controlled gene *daywake*. There was no overlap in significant, downregulated genes expressed at 7 DPP in *H. erato*. At 5 DPP in *H. erato*, there were no significant, concordantly DE genes. However, a *doublesex-like* gene on chromosome 8 narrowly missed the significance cutoff and was downregulated (LogFC < −1.5) in both comparisons (FDR = 0.02 between subspecies, FDR = 0.08 between wing regions). In *H. melpomene* at both 7 and 5 DPP there was no overlap between genes that were DE between subspecies and wing regions.

Genes involved in controlling scale structure may be similarly differentially expressed between species. Between subspecies, at 7 DPP there were no concordantly DE genes in either species. However, at 5 DPP, there were 2 concordant genes significantly DE, *Fatty acid synthase* and *Gamma-glutamylcyclotransferase* (Table S14). For the wing region comparison, at 7 DPP there were 4 concordant genes significantly DE in both species, the homeobox gene *invected, Transglutaminase, uncharacterized LOC113401078* and the *doublesex-like* gene, which was also DE between *H. erato* subspecies (at 5 DPP), but none at 5 DPP (although the *doublesex*-like gene is again DE in *H. melpomene*, Table S14).

### DE genes in the QTL intervals

In order to identify candidate genes in the QTL intervals, we identified DE genes within these genomic regions. In *H. erato*, there were 2 and 5 DE genes in the ‘BR’ interval on chromosome 20 at 5 DPP and 7 DPP, respectively (Table S15). One of the genes at 7 DPP was *Fringe*, a boundary specific signalling molecule which modulates the Notch signalling pathway and has roles in eyespot formation and scale cell spacing in butterflies (Reed and Serfas, 2004; Thurmond *et al*., 2018).

On the Z chromosome, at 5 DPP there were 27, 25, and 17 genes significantly DE between subspecies in the ‘ridge spacing’, ‘luminance’ and ‘BR’ intervals, respectively, with 16 genes shared between all 3 intervals (Figure 4, Table S15). Of note, the microtubule motor protein, *dynein heavy chain 6* was within all three QTL intervals and highly-upregulated (LogFC > 3.0, FDR < 0.05) in the iridescent subspecies. Additionally, an *O-GlcNAc transferase*, with strong similarity to *Drosophila* polycomb group gene *super sex combs* was highly differentially expressed (LogFC = −9.32, FDR<0.004) and matched the exact physical location of the ‘BR’ and ‘luminance’ marker within the genome.

**Figure 4.**
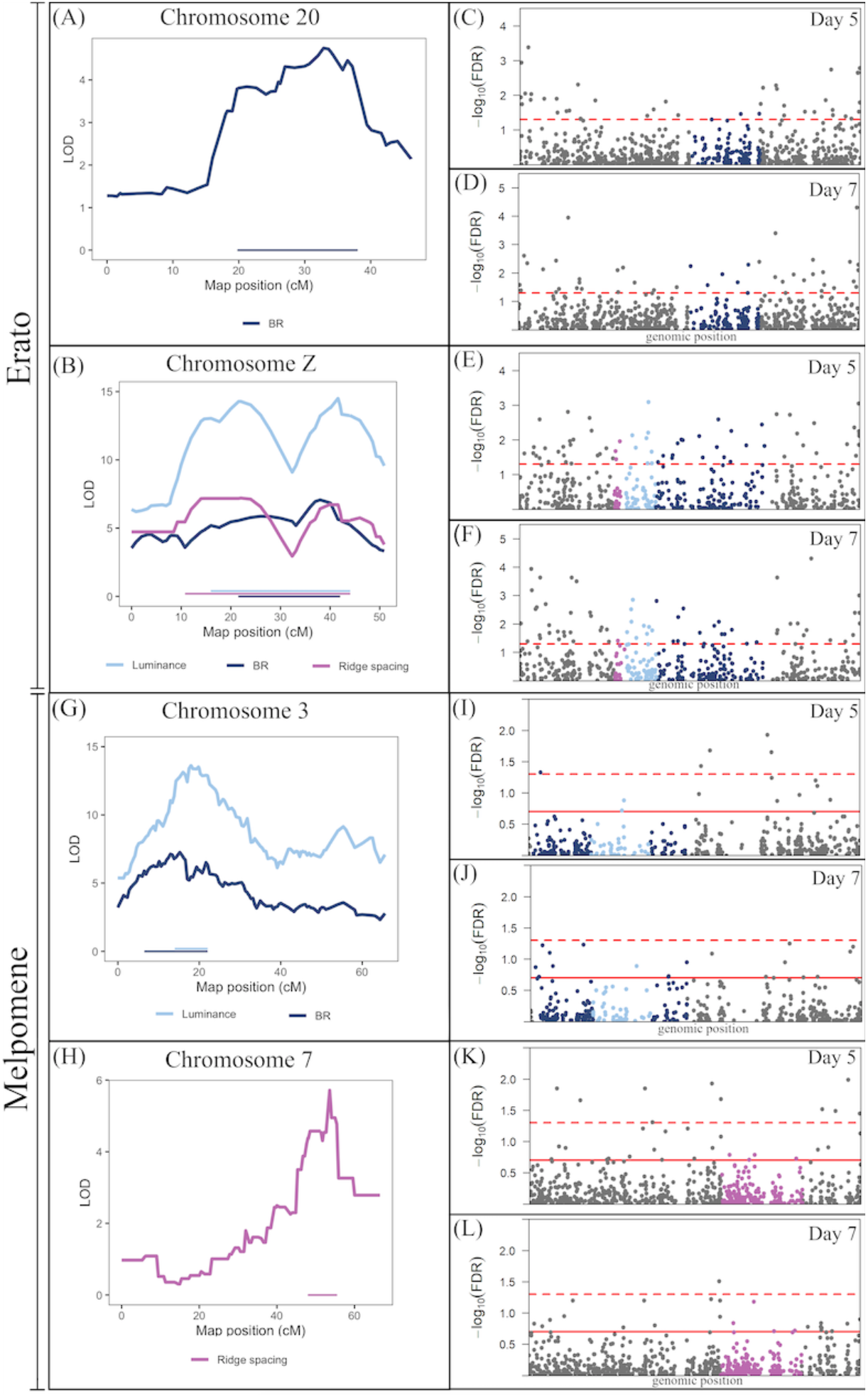
Differential expression of genes in the QTL in *Heliconius erato* (A-F) and *Heliconius melpomene* (G-L). Left panels: LOD scores and QTL intervals in *H. erato* (A, B) and *H. melpomene* (G, H). Right panels: -log10 False Discovery Rate (FDR) for differential expression. Genes are coloured within the QTL intervals for *H. erato* (C-F) and *H. melpomene* (I-L), with colours matching those of the intervals in the panels on the left. In E and F the QTL overlap, such that all genes in the BR and ridge spacing intervals also fall within the luminance interval, see Table S15 for details. In I and J the luminance interval is within the BR interval. The dashed red line indicates FDR = 0.05 (significance), solid red line indicates FDR = 0.2.

At 7 DPP, on the Z chromosome there were 24, 23, and 14 genes significantly DE in the ‘ridge spacing’, ‘luminance’ and ‘BR’ intervals, respectively, with 14 shared across all 3 regions (Table S15). The gene *trio*, which functions in actin structure regulation through activation of Rho-family GTPases (Thurmond *et al*., 2018), was found in all three intervals with particular proximity to the ‘ridge spacing’ marker (405 kbp away from the start of this gene). In addition to the functional role of *trio*, its high expression and large fold change (logCPM = 7.34, LogFC = −2.29, FDR = 0.0015) makes it a particularly good candidate for a role in optical nanostructure development in *H. erato*. Furthermore, a novel gene (MSTRG.21985) was also DE expressed (LogFC= −1.28, FDR = 0.0115) and may be part of a Rho GTPase activating protein (182 bp upstream of a gene with this annotation).

In *H. melpomene*, there were no DE genes between subspecies in the ‘ridge spacing’ interval on chromosome 7 at either stage. However, 7 DPP, the gene *ringmaker*, which functions in microtubule organisation (Thurmond *et al*., 2018) showed slight DE (logFC = −1.43, FDR = 0.144). On chromosome 3, in the BR interval there was 1 novel gene (MSTRG.3173) DE at 5 DPP (but this falls outside the luminance interval) and no DE genes at 7 DPP (Table S16). The gene *miniature*, which in fly bristles is a component of the cuticulin envelope functioning in interactions between the depositing cuticle, membrane and cytoskeleton (Roch, Alonso and Akam, 2003), falls in the overlap of the luminance and BR regions and shows slight DE at 5 DPP (logFC = 1.60, FDR = 0.192).

For the wing region comparison, in *H. erato* there were no genes DE at either stage within any of the QTL intervals. For *H. melpomene*, there was 1 DE gene in the ‘BR’ interval (but outside the ‘luminance’ interval) on chromosome 3 at 7 DPP (a *lactase-phlorizin hydrolase-like* gene) and no DE genes at 5 DPP. For the ‘Ridge Spacing’ interval on chromosome 7 there was 1 DE gene at 5 DPP, an *F-actin-uncapping protein LRRC16A* and 1 gene at 7 DPP, a *cuticle protein 18.6-like* gene (Table S17).

## Discussion

In one of the first studies to look at the genetics of structural colour variation in terms of both colour and structure, we show that the trait is controlled by multiple genes in the co-mimics *Heliconius erato* and *Heliconius melpomene*. While we found only a small number of QTL, these explain relatively little of the overall phenotypic variation, suggesting there are more loci which remain undetected. Some of these may be the genes that we detected as differentially expressed, but that fall outside the detected QTL intervals. Of particular interest are genes that we detected as differentially expressed both between subspecies and between wing regions that differ in scale type. *Chitin deacetylase 1* is one such candidate in *H. erato*, which is on chromosome 5 (not in a QTL interval). Chitin is the main component of the cuticle and the differential expression of a potential chitin-degrading gene could alter the formation of the scale ridges (Yu *et al*., 2016).

Within each species, we find that hue and brightness (BR and luminance) are controlled by loci on the same chromosomes. In *H. erato*, this was on the Z chromosome, confirming our previous phenotypic analysis (Brien *et al*., 2018), and in *H. melpomene*, on chromosome 3. An additional locus on chromosome 20 was also found to affect blue colour but not brightness in *H. erato*. The Z chromosome locus in *H. erato* appears to control ridge spacing, which could have a direct effect on the brightness of the reflectance by increasing the density of reflective structures. Indeed, in the single-family analyses, luminance and ridge spacing mapped to exactly the same marker. However, the observed correlation between brightness and ridge spacing in *H. erato* may be a product of an unobserved association between tighter ridge spacing and other aspects of scale nanostructure, specifically the number of lamellae layers within the ridges. Theoretical analyses and simulations of the optical properties of multilayers have revealed that increasing the number of layers will result in a rapid increase of brightness; adding even a small number of layers produces a significant increase in the amount of reflected light (Kinoshita and Yoshioka, 2005). Therefore, the Z chromosome locus may be affecting multiple aspects of scale structure, producing the observed correlations between the different colour and structure measurements. Indeed, some DE genes in the Z locus may control multiple aspects of scale structure. For example, *trio* acts in several signalling pathways to promote reorganisation of the actin cytoskeleton through Rho GTPase activation. Its regulatory function may be repeatedly employed during scale development in the formation of different aspects of scale ultrastructure guided by the actin cytoskeleton. Interestingly, potentially related signalling genes, such as the novel gene located immediately before a Rho GTPase activating protein, also fall within this locus and are DE, potentially suggesting there are several functional genes linked together in this region.

In contrast, in *H. melpomene* we found different loci controlling colour and ridge spacing, suggesting a more dispersed genetic architecture and different loci controlling different aspects of scale structure. We found strong evidence for a locus on chromosome 3 controlling BR and luminance, but this locus appeared to have no effect on our measurements of scale structure and so is likely controlling other aspects of scale structure not quantified here. Instead, we find a locus on chromosome 7 that partially controls ridge spacing. We see a small, but not significant, effect of this chromosome on BR colour, suggesting that ridge spacing may have a small and relatively weak effect on colour in *H. melpomene*, despite the parental populations showing a similar difference in ridge spacing to that seen in *H. erato*. If *H. erato* has a locus on the Z chromosome that can control multiple aspects of scale structure, while *H. melpomene* requires mutations at loci dispersed around the genome, this could provide one explanation for how *H. erato* has been able to evolve brighter structural colour than that observed in *H. melpomene*.

In contrast to many of the loci for pigment colour patterns which are homologous across multiple *Heliconius* species, the loci controlling iridescence in *H. erato* and *H. melpomene* appear to be largely different. Differences in the physical scale architecture and brightness of colour between the species perhaps makes these genetic differences unsurprising (Parnell *et al*., 2018; Curran *et al*., 2020). A lack of genetic parallelism may also be more likely for a quantitative trait such as iridescence (Conte *et al*., 2012). Nevertheless, on the Z chromosome in *H. melpomene*, we do observe elevated LOD scores in the QTL analysis and low p-value SNPs in the GWAS for both scale structure traits, but neither of the colour traits. This suggests that *H. melpomene* may have a locus homologous to that in *H. erato*, which is controlling some aspects of scale structure variation, but with apparently little or no effect on colour variation. In addition, we find some genes that appear to show parallel expression patterns between species. Of particular interest is a *doublesex-like* gene that is DE between wing regions in both species and between *H. erato* subspecies. A different duplication of *doublesex* has been found to control structural colour in the Dogface butterfly (Rodriguez-Caro *et al*., 2021), making this an interesting, potentially parallel candidate between species. It is possible that the evolutionary pathways may be different between species, but have triggered expression changes in similar downstream developmental pathways. However, we found very few genes that show concordant expression patterns between species.

In recent years, reverse genetics research has revealed a surprising connection between the molecular machinery underlying the development of pigmented wing patterns and the ultrastructure of butterfly scales in various species (Zhang, Mazo-Vargas and Reed, 2017; Concha *et al*., 2019; Fenner *et al*., 2020; Livraghi *et al*., 2021). However, our QTL are not associated with any known colour pattern gene of large or small effect in *Heliconius (aristaless, WntA, vvl, cortex* and *optix* - located on chromosomes 1, 10, 13, 15 and 18 respectively) (Nadeau, 2016). Our findings show that *H. erato* and *H. melpomene* do not use the known molecular machinery of wing pattern production for sculpting specialised nanostructures and iridescent wings, and that the production of structural colour is completely decoupled from that of mimicry related wing pattern regulation and pigment production.

Overall, we show major differences in the genetic basis of structural colour in *H. erato* and *H. melpomene*. Combining this with gene expression analyses, we have been able to identify novel candidate genes for the control of structural colour variation with potential functions in chitin metabolism, cytoskeleton formation, gene expression regulation and cell signalling.

## Acknowledgements

We thank the governments of Ecuador and Panama for permission to collect butterflies. Thanks to Darwin Chalá, Juan López and Gabriela Irazábal for their assistance with the crosses. We are grateful to the European Synchrotron Radiation Facility for provision of X-ray beamtime under proposal LS2720, and to Andrew Dennison for assistance with USAXS data collection. We also thank Alexandre Thiery for his assistance with the differential expression analysis. Library preparation and sequencing was performed by staff at the Edinburgh Genomics Facility, at the University of Edinburgh. We thank Alan Dunbar for use of the scanning electron microscope.

## Data availability

Genomic sequence data from the crosses are deposited in the European Nucleotide Archive under project number PRJEB38330. RNA sequence data are deposited under project number XXXX Photographs of all samples can be found at https://doi.org/10.5281/zenodo.3799188. USAXS data handling scripts can be found at https://github.com/juanenciso14/butterfly_usaxs. Linkage maps, QTL scripts and all phenotypic measurements used will be available on publication.

## Author contributions

MNB and JER performed the QTL analyses and wrote the manuscript with NJN and VJL. PR and MNB constructed the linkage maps. AJP co-ordinated collection of the USAXS data, which were collected by TZ, AJP, EVC, NJN and MNB. JER analysed the USAXS and SEM data and ran the association analysis. PAS, CM, MNB, EVC and NJN performed the crosses. NJN collected the tissue and extracted the RNA for the gene expression analysis, VJL performed all gene expression analyses. NJN devised and co-ordinated the study. All authors read and commented on the manuscript.

