## Supplementary material for "The genetic basis of structural colour variation in mimetic *Heliconius* butterflies": Figure S1

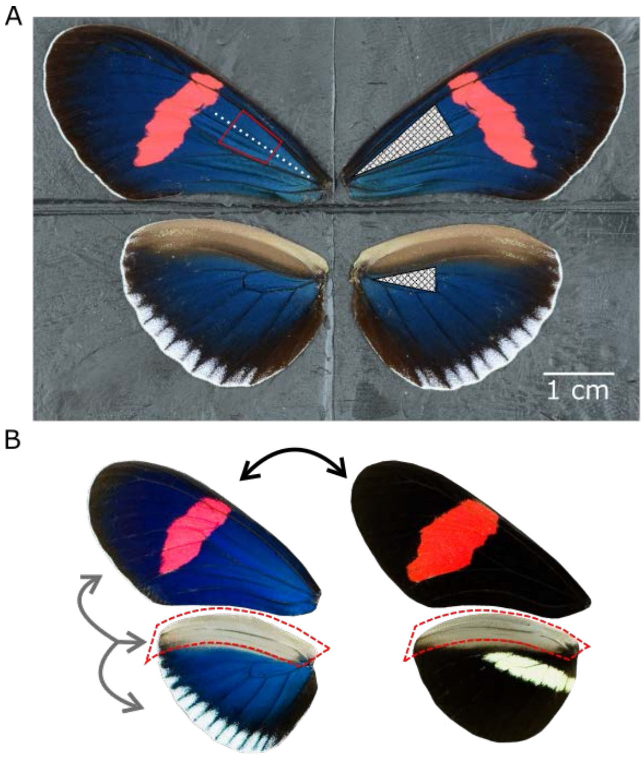


Figure S1: (A) Wing regions measured for colour analysis (hatched triangles on right side); for USAXS experiment (white dashed line on left forewing); and wing region cut for SEM imaging (red box on left forewing). (B) Wing regions and comparisons used for RNA sequencing and gene expression analysis. The red dashed box is the androconial region, which was dissected and sequenced separately from the rest of the fore- and hind-wings (which were pooled). Arrows depict the two main comparisons: Black arrow, between subspecies comparison, in which the non-iridescent (Panama) subspecies is compared to the iridescent (Ecuador) subspecies (for the main parts of the wing); grey arrow is the wing region comparison in which the iridescent wing regions are compared to the androconial wing region (for the iridescent subspecies). Image in A is one of the actual images (of *H. melpomene*) used in the colour analysis. Images in B are for illustrative purposes only (depicting adult *H. erato* wings). The actual dissected wings were from developing pupae and had no colour. Wing veins were used as landmarks for dissecting the androconial region (with the cut made along the Rs or vein 7, the second most anterior hindwing vein).

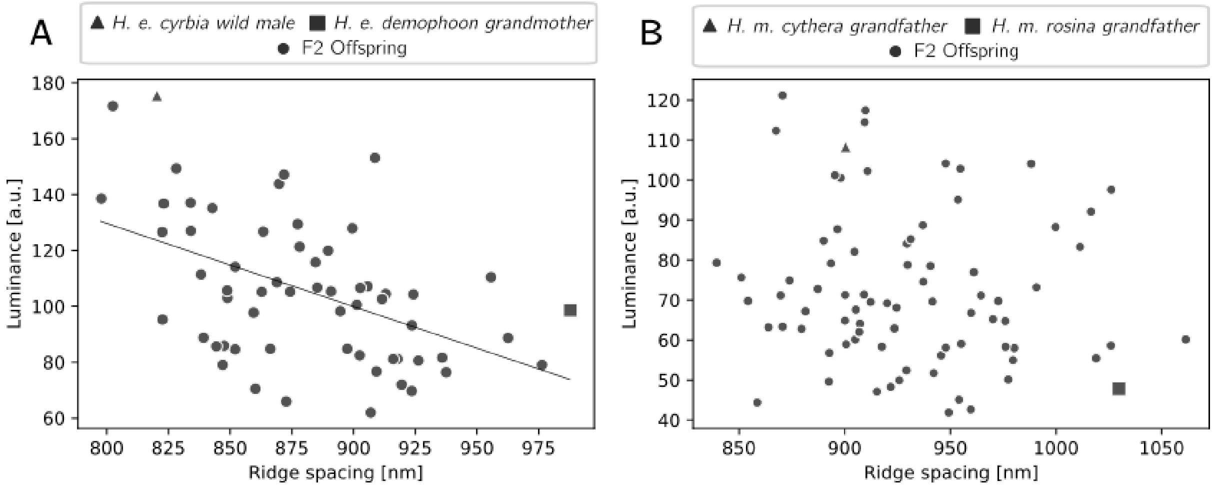


Figure S2. Relationship between ridge spacing and luminance in F2 offspring of *H. erato* (A) and *H. melpomene* (B). A significant correlation was found for *H. erato* only.

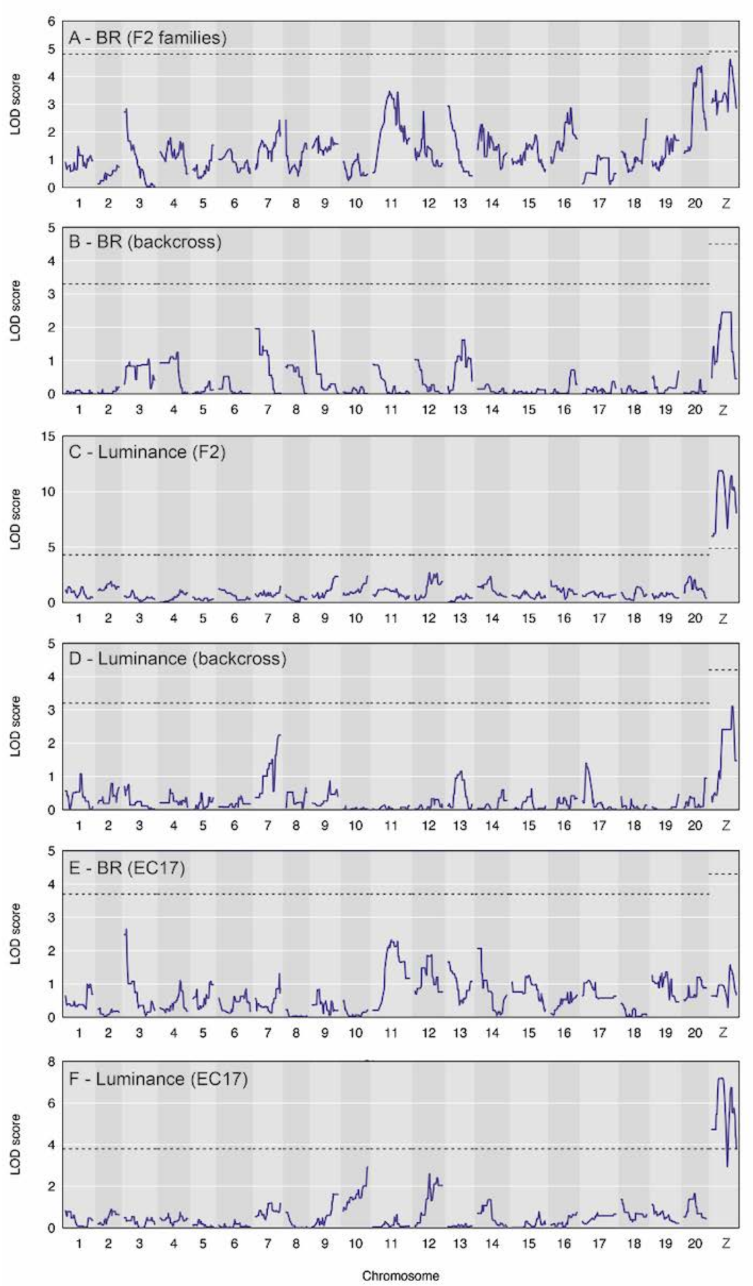


Figure S3: QTL plots for *H. erato* F2 families combined, the backcross family, and the family for which we have scale structure data (EC17) separately.

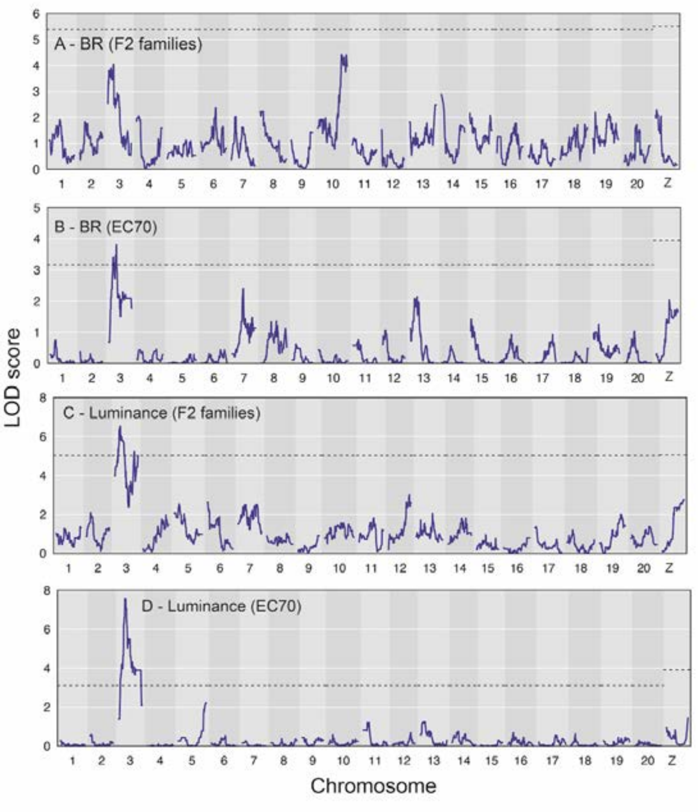


Figure S4: QTL plots for *H. melpomene* (A, C) the three F2 families combined and (B, D) the EC70 brood.

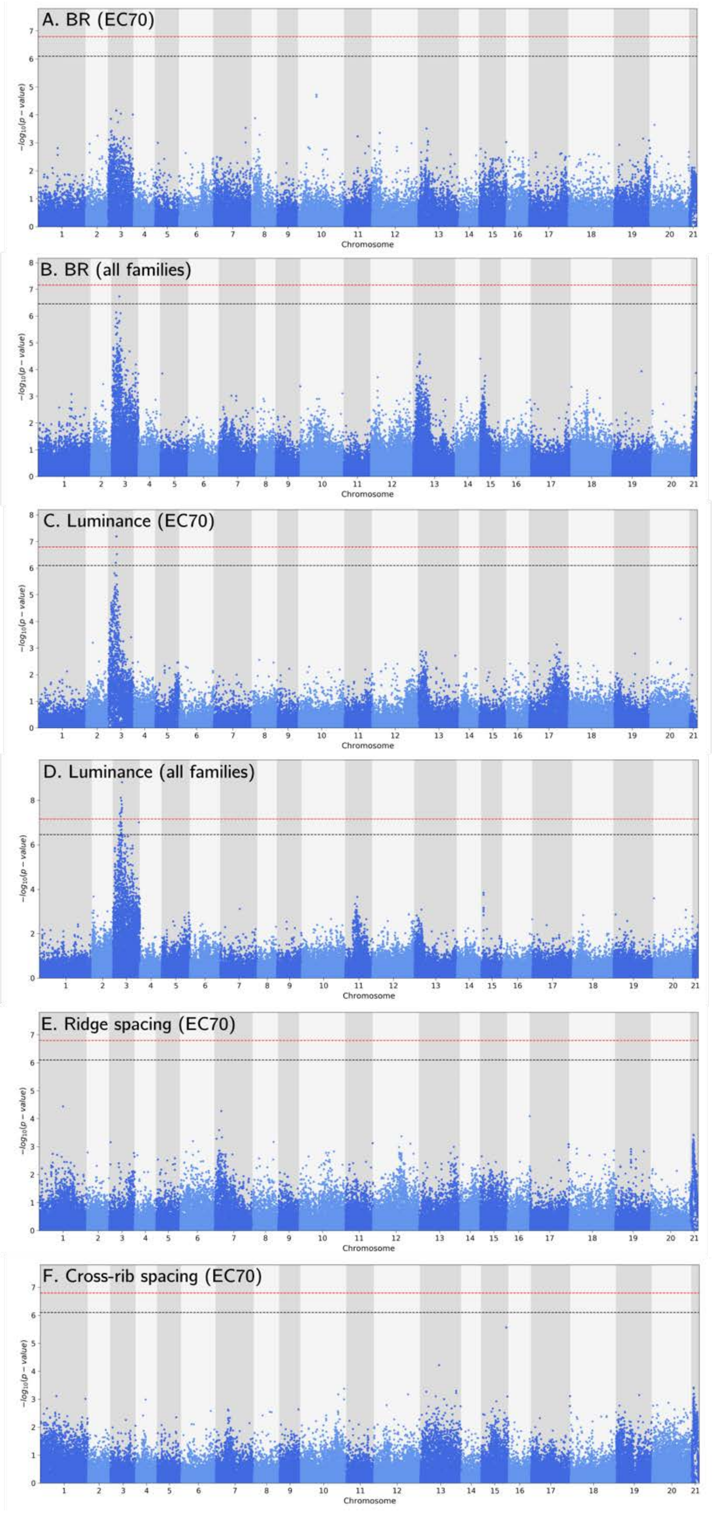


Figure S5: Association analysis for four phenotypes in *H. melpomene*. p = 0.05 significance: black dashed line, p = 0.01: red dashed line.

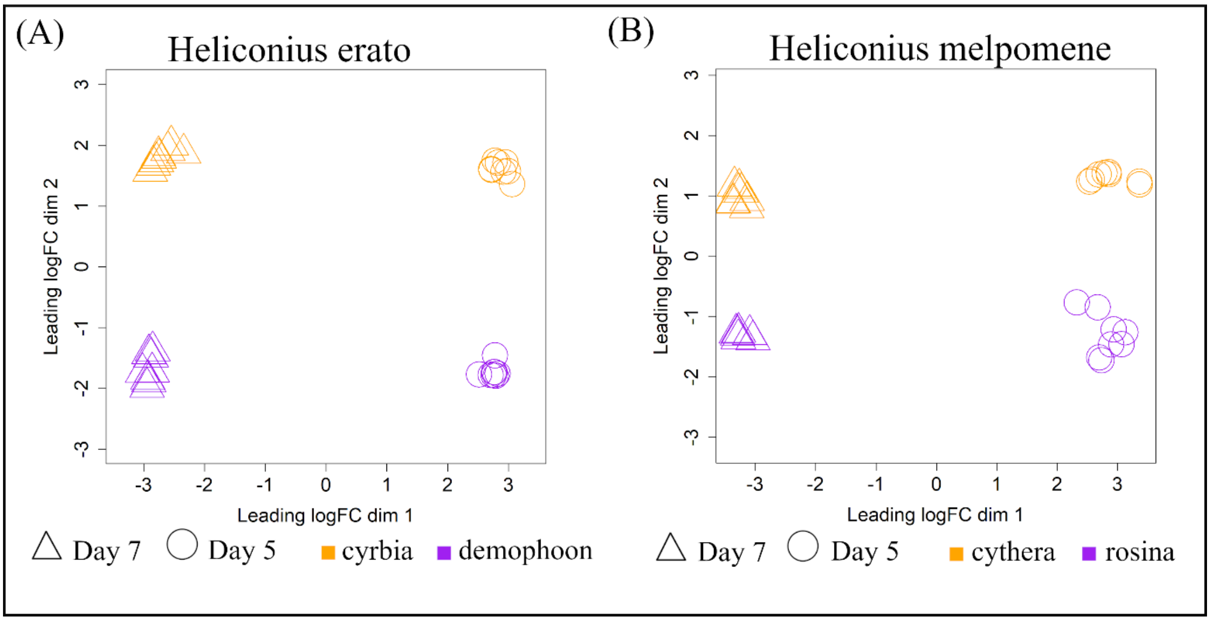


Figure S6. Multidimensional scaling of differences in expression profiles between samples of (A) *Heliconius erato* and (B) *Heliconius melpomene*. Clustering of samples is based on filtered and normalised expression levels. Point shape indicates stage and colour indicates race.

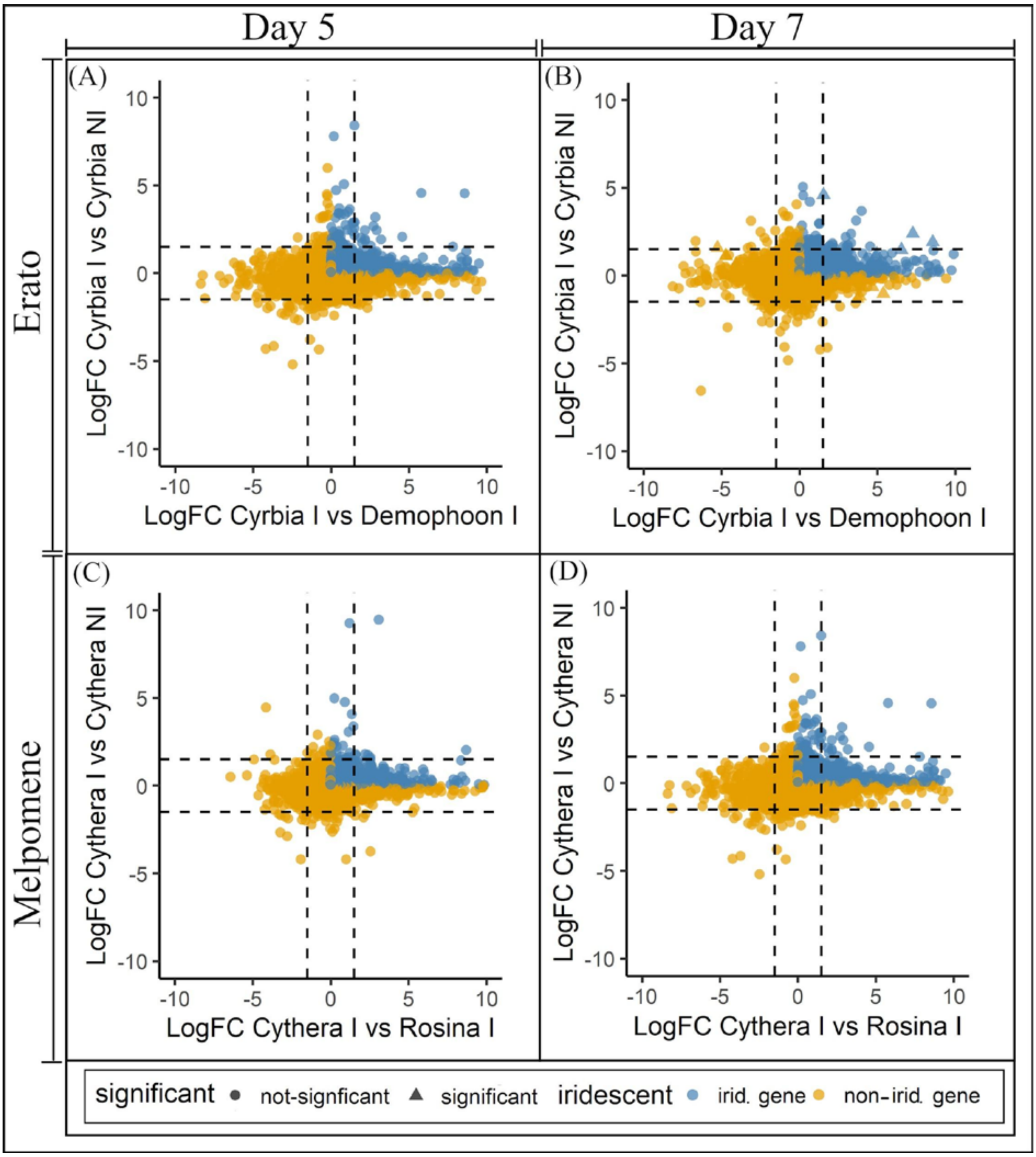


Figure S7: Overlapping expression patterns of differentially expressed genes in race and tissue comparisons of *H. erato* (A-B) and *H. melpomene* (C-D) at 5 and 7 DPP. Genes which are upregulated (Log FC > 0) in both the race comparison (iridescent race vs non-iridescent race) and the wing region comparison (iridescent region vs non-iridescent region of iridescent race) are likely iridescent genes (blue points). Symbols indicate whether a gene falls within the significance level cut-off (FDR < 0.05) for both the race and tissue comparison. Dashed lines indicate LogFC < -1.5 and LogFC > 1.5 and therefore indicate highly expressed genes.

Table S1: Details of crosses and offspring used for phenotyping colour and scale structure, and those sequenced to produce the linkage maps. Parents of crosses have the cross they originated from in brackets.

| **Cross ID** | **Cross type** | **Father ID** | **Mother ID** | **Offspring phenotyped for:** | | **Sequenced to produce l[illegible] maps** |
|---|---|---|---|---|---|---|
| | | | | BR/luminance | Scale structures | |
| ***Heliconius erato*** | | | | | | |
| EC01F1 | *demophoon ♂x cyrbia ♀* | 14N012 | 14N011 | | | 4 |
| EC10F1 | *cyrbia ♂ x demophoon ♀* | 14N065 | 14N064 | | | 3 |
| EC39F1 | *cyrbia ♂ x demophoon ♀* | 14N339 | 14N338 | | | 2 |
| EC45F1 | *demophoon ♂x cyrbia ♀* | 14N359 | 14N358 | | | 2 |
| EC41BC | *cyrbia ♀ x (cyrbia ♂ x demophoon ♀)* | 14N348 (EC10) | 14N347 | 40 | | 40 |
| EC13F2 | *demophoon maternal grandfather* | 14N078 (EC01) | 14N077 (EC01) | 3 | | 3 |
| EC15F2 | *demophoon maternal grandfather* | 14N093 (EC01) | 14N089 (EC01) | 5 | | 5 |
| EC17F2 | *cyrbia maternal grandfather* | 14N112 (EC01) | 14N111 (EC10) | 56 | 56 | 56 |
| EC18F2 | *cyrbia maternal grandfather* | 14N114 (EC01) | 14N113 (EC10) | 14 | | 14 |
| EC53F2 | *cyrbia maternal grandfather* | 14N396 (EC39) | 14N395 (EC39) | 21 | | 21 |
| **Total** | | | | | | 150 |
| ***Heliconius melpomene*** | | | | | | |
| EC05F1 | *cythera ♂x rosina ♀* | 14N052 | 14N051 | | | |
| EC07F1 | *cythera ♂x rosina ♀* | 14N059 | 14N058 | | | |
| EC09F1 | *rosina ♂ x cythera ♀* | 14N063 | 14N062 | | | |
| EC26F1 | *cythera ♂x rosina ♀* | 14N219 | 14N220 | | | |
| EC27F1 | *rosina ♂ x cythera ♀* | 14N221 | 14N222 | | | |
| EC35F1 | *cythera ♂x rosina ♀* | Unknown *cythera* | 14N315 | | | |
| EC38F1 | *rosina ♂ x cythera ♀* | 14N337 | 14N336 | | | |
| EC48F1 | *rosina ♂ x cythera ♀* | 14N366 | Unknown *cythera* | | | 2 |
| EC49F1 | *cythera ♂x rosina ♀* | 14N368 | 14N367 | | | 3 |
| EC51F2 | *rosina maternal grandfather* | 14N379 (EC35) | 14N385 (EC38) | | | |

| | | | | | | |
|---|---|---|---|---|---|---|
| EC57F2 | *rosina maternal grandfather* | 14N452 (EC38) | 14N451 (EC48) | | | |
| EC63F2 | *cythera maternal grandfather* | 14N480 (EC48) | 14N479 (EC49) | 52 | | 52 |
| EC65F2 | *cythera maternal grandfather* | 15N563 (EC48) | 15N562 (EC49) | 54 | | 54 |
| EC69F2 | *rosina maternal grandfather* | 15N605 (EC49) | 15N604 (EC48) | 7 | | 7 |
| EC70 | Unknown maternal grandfather | 15N614 (EC49) | 15N613 | 106 | 73 | 106 |
| **Total** | | | | | | 224 |

Table S2: Samples of *H. erato* and *H. melpomene* used in the RNAseq analysis.
* indicates the *H. melpomene* sample which was removed from further DE analyses.

| **Sample** | **Subspecies** | **Stage** | **Tissue** | **Individual** | **Sex** |
|---|---|---|---|---|---|
| ***H. erato*** | | | | | |
| M_23I | cyrbia | 5 | FW + post-HW | 1 | Male |
| M_23N | cyrbia | 5 | anterior-HW | 1 | Male |
| F_27I | cyrbia | 5 | FW + post-HW | 2 | Female |
| F_27N | cyrbia | 5 | anterior-HW | 2 | Female |
| F_25I | cyrbia | 5 | FW + post-HW | 3 | Female |
| F_25N | cyrbia | 5 | anterior-HW | 3 | Female |
| M_28I | cyrbia | 5 | FW + post-HW | 4 | Male |
| M_28N | cyrbia | 5 | anterior-HW | 4 | Male |
| F_31I | demophoon | 5 | FW + post-HW | 1 | Female |
| F_31N | demophoon | 5 | anterior-HW | 1 | Female |
| F_32I | demophoon | 5 | FW + post-HW | 2 | Female |
| F_32N | demophoon | 5 | anterior-HW | 2 | Female |
| F_20I | demophoon | 5 | FW + post-HW | 3 | Female |
| F_20N | demophoon | 5 | anterior-HW | 3 | Female |
| M_36I | demophoon | 5 | FW + post-HW | 4 | Male |
| M_36N | demophoon | 5 | anterior-HW | 4 | Male |
| M_43I | cyrbia | 7 | FW + post-HW | 1 | Male |
| M_43N | cyrbia | 7 | anterior-HW | 1 | Male |
| M_44I | cyrbia | 7 | FW + post-HW | 2 | Male |
| M_44N | cyrbia | 7 | anterior-HW | 2 | Male |
| M_50I | cyrbia | 7 | FW + post-HW | 3 | Male |
| M_50N | cyrbia | 7 | anterior-HW | 3 | Male |
| M_51I | cyrbia | 7 | FW + post-HW | 4 | Male |
| M_51N | cyrbia | 7 | anterior-HW | 4 | Male |
| F_39I | demophoon | 7 | FW + post-HW | 1 | Female |
| F_39N | demophoon | 7 | anterior-HW | 1 | Female |
| F_40I | demophoon | 7 | FW + post-HW | 2 | Female |

| | | | | | |
|---|---|---|---|---|---|
| F_40N | demophoon | 7 | anterior-HW | 2 | Female |
| M_45I | demophoon | 7 | FW + post-HW | 3 | Male |
| M_45N | demophoon | 7 | anterior-HW | 3 | Male |
| F_47I | demophoon | 7 | FW + post-HW | 4 | Female |
| F_47N | demophoon | 7 | anterior-HW | 4 | Female |
| ***H. melpomene*** | | | | | |
| S_84AF | cythera | 5 | anterior-HW | 1 | Female |
| S_84IF | cythera | 5 | FW + post-HW | 1 | Female |
| S_80AM | cythera | 5 | anterior-HW | 2 | Male |
| S_80IM | cythera | 5 | FW + post-HW | 2 | Male |
| S_91AF | cythera | 5 | anterior-HW | 3 | Female |
| S_91IF | cythera | 5 | FW + post-HW | 3 | Female |
| S_81AM | cythera | 5 | anterior-HW | 4 | Male |
| S_81IM | cythera | 5 | FW + post-HW | 4 | Male |
| S_62AM | rosina | 5 | anterior-HW | 1 | Male |
| S_62IM | rosina | 5 | FW + post-HW | 1 | Male |
| S_76AF | rosina | 5 | anterior-HW | 2 | Female |
| S_76IF | rosina | 5 | FW + post-HW | 2 | Female |
| S_78AM | rosina | 5 | anterior-HW | 3 | Male |
| S_78IM | rosina | 5 | FW + post-HW | 3 | Male |
| S_90AF | rosina | 5 | anterior-HW | 4 | Female |
| S_90IF | rosina | 5 | FW + post-HW | 4 | Female |
| S_71AM | cythera | 7 | anterior-HW | 1 | Male |
| S_71IM | cythera | 7 | FW + post-HW | 1 | Male |
| S_96AF | cythera | 7 | anterior-HW | 2 | Female |
| S_96IF | cythera | 7 | FW + post-HW | 2 | Female |
| S_86AM | cythera | 7 | anterior-HW | 3 | Male |
| S_86IM | cythera | 7 | FW + post-HW | 3 | Male |
| S_61AF | cythera | 7 | anterior-HW | 4 | Female |
| S_61IF | cythera | 7 | FW + post-HW | 4 | Female |
| S_60AM* | rosina | 7 | anterior-HW | 1 | Male |

| | | | | | |
|---|---|---|---|---|---|
| S_60IM* | rosina | 7 | FW + post-HW | 1 | Male |
| S_65AM | rosina | 7 | anterior-HW | 2 | Male |
| S_65IM | rosina | 7 | FW + post-HW | 2 | Male |
| S_85AM | rosina | 7 | anterior-HW | 3 | Male |
| S_85IM | rosina | 7 | FW + post-HW | 3 | Male |
| S_74AM | rosina | 7 | anterior-HW | 4 | Male |
| S_74IM | rosina | 7 | FW + post-HW | 4 | Male |

Table S3: Markers with the highest LOD scores in each QTL analysis for *H. erato*, using separate F2 and backcross families, and all families combined. Variance is not calculated for the combined scans.

| **Phenotype** | **Families** | **Chr** | **Marker (LG_snpposition)** | **Position (cM)** | **LOD** | **p** | **95% confidence interval lower** | **interval upper** | **% variance explained** |
|---|---|---|---|---|---|---|---|---|---|
| BR | All combined | Z | Herato2101_12449252 | 38 | 7.07 | 0.001 | Herato2101_7003910 | Herato2101_12672960 | |
| BR | All combined | 20 | Herato2001_12633065 | 32.9 | 4.75 | 0.022 | Herato2001_9722974 | Herato2001_13728343 | |
| Luminance | All combined | Z | Herato2101_12449398 | 41.6 | 14.5 | <0.001 | Herato2101_5394360 | Herato2101_12673089 | |
| Ridge Spacing | EC17 | Z | Herato2101_7491127 | 23 | 5.21 | 0.013 | Herato2101_4831239 | Herato2101_12449360 | 34.9 |
| BR | EC17 | | None | | | | | | |
| Luminance | EC17 | Z | Herato2101_7491127 | 22 | 7.2 | <0.001 | Herato2101_5394170 | Herato2101_12449439 | 44.8 |
| BR | F2 | 20 | Herato2001_13728260 | 36.5 | 4.38 | NS 0.097 | Herato2001_11442706 | Herato2001_13728343 | 12.3 |
| BR | F2 | Z | Herato2101_12449252 | 38 | 4.62 | NS 0.143 | Herato2101_3280129 | Herato2101_12673089 | 19.5 |
| Luminance | F2 | Z | Herato2101_7491127 | 22 | 11.9 | <0.001 | Herato2101_5394170 | Herato2101_7491127 | 40.2 |
| BR | Backcross | | None | | | | | | |
| Luminance | Backcross | | None | | | | | | |

Table S4: Markers with the highest LOD scores in each QTL analysis for *H. melpomene*, using F2 families and the EC70 brood separately, and all families combined.

| **Phenotype** | **Families** | **Chr** | **Marker (LG_snpposition)** | **Position (cM)** | **LOD** | **p** | **95% confidence interval lower** | **interval upper** | **% variance** |
|---|---|---|---|---|---|---|---|---|---|
| BR | All combined | 3 | Hmel203003o_2119654 | 15.22 | 7.26 | 0.001 | Hmel203003o_121866 | Hmel203003o_4953744 | |
| Luminance | All combined | 3 | Hmel203003o_2635435 | 17.97 | 13.61 | <0.001 | Hmel203003o_1984108 | Hmel203003o_3795412 | |
| Ridge spacing | EC70 | 7 | Hmel207001o_11550301 | 53.61 | 5.71 | <0.001 | Hmel207001o_8304361 | Hmel207001o_11770725 | 30.3 |
| BR | EC70 | 3 | Hmel203003o_3638237 | 21.6 | 3.82 | 0.021 | Hmel203003o_775817 | Hmel203003o_9744554 | 15.29 |
| BR | F2 | 3 | Hmel203003o_2119869 | 16.14 | 4.03 | NS | Hmel203003o_121866 | Hmel203003o_4953744 | 9.16 |
| BR | F2 | 10 | Hmel210001o_15604785 | 66.83 | 4.41 | NS | Hmel210001o_15497435 | Hmel210001o_17837389 | 16.73 |
| Luminance | EC70 | 3 | Hmel203003o_2647602 | 18.32 | 7.57 | <0.001 | Hmel203003o_2647602 | Hmel203003o_4028331 | 28 |
| Luminance | F2 | 3 | Hmel203003o_2119654 | 15.22 | 6.52 | 0.0042 | Hmel203003o_1264960 | Hmel203003o_9987035 | 9.43 |

Table S5: Genes differentially expressed (FDR<0.2) between *H. erato cyrbia* and *H. erato demophoon* at 5 days post-pupation (DPP) - see Excel spreadsheet

Table S6: Genes differentially expressed (FDR<0.2) between *H. erato cyrbia* and *H. erato demophoon* at 7 DPP. - see Excel spreadsheet

Table S7: Genes differentially expressed (FDR<0.2) between *H. melpomene cythera* and *H. melpomene rosina* at 5 DPP - see Excel spreadsheet

Table S8: Genes differentially expressed (FDR<0.2) between *H. melpomene cythera* and *H. melpomene rosina* at 7 DPP - see Excel spreadsheet

Table S9: Genes differentially expressed (FDR<0.2) between wing regions in *H. erato cyrbia* at 5 DPP - see Excel spreadsheet

Table S10: Genes differentially expressed (FDR<0.2) between wing regions in *H. erato cyrbia* at 7 DPP. - see Excel spreadsheet

Table S11: Genes differentially expressed (FDR<0.2) between wing regions in *H. melpomene cythera* at 5 DPP - see Excel spreadsheet

Table S12: Genes differentially expressed (FDR<0.2) between wing regions in *H. melpomene cythera* at 7 DPP - see Excel spreadsheet

Table S13. Overlapping genes (FDR < 0.2) between the wing region comparison of the iridescent subspecies and the subspecies comparison. Bold indicates significant genes with an FDR < 0.05 in both comparisons.

| gene | logFC.race | logCPM.race | F.race | FDR.race | logFC.tissue | logCPM.tissue | F.tissue | FDR.tissue | Annotation | E-value |
|---|---|---|---|---|---|---|---|---|---|---|
| ***Heliconius erato*** **DAY 5** | | | | | | | | | | |
| **MSTRG.13758\|evm.TU.Herato1301.635** | 12.664 | 3.502 | 39.655 | 0.020 | 1.280 | 3.995 | 50.369 | 0.179 | U8-agatoxin-Ao1a-like [Vanessa tameamea] | 4.00E-66 |
| **MSTRG.7192\|evm.TU.Herato0801.87** | -2.719 | 4.719 | 24.967 | 0.019 | -1.839 | 3.853 | 91.115 | 0.080 | protein doublesex-like isoform X1 [Vanessa tameamea] | 2.00E-29 |
| **MSTRG.13445\|evm.TU.Herato1301.472** | -0.903 | 3.814 | 6.293 | 0.194 | -0.974 | 3.843 | 47.992 | 0.179 | transcription factor cwo isoform X4 [Vanessa tameamea] | 0 |
| **DAY 7** | | | | | | | | | | |
| **MSTRG.22333** | 5.601 | 4.269 | 278.480 | 0.000 | 0.370 | 5.065 | 16.352 | 0.154 | na | na |
| **MSTRG.21653\|evm.TU.Herato2101.110** | 3.634 | 3.755 | 160.991 | 0.000 | 0.741 | 4.324 | 28.989 | 0.084 | probable cytochrome P450 303a1 isoform X2 [Danaus plexippus plexippus] | 0 |
| **MSTRG.10205\|evm.TU.Herato1007.170** | 4.283 | 7.069 | 90.206 | 0.002 | 0.379 | 7.812 | 14.481 | 0.175 | carboxypeptidase N subunit 2-like [Vanessa tameamea] | 0 |

| | | | | | | | | | | |
|---|---|---|---|---|---|---|---|---|---|---|
| **MSTRG.18455\|evm.TU.Herato1805.101\|evm.TU.Herato1805.102** | 1.983 | 4.676 | 46.658 | 0.006 | 1.174 | 4.893 | 17.173 | 0.187 | circadian clock-controlled protein-like [Vanessa tameamea] | 7.00E-100 |
| **MSTRG.13019** | 6.520 | -0.145 | 36.588 | 0.010 | 1.480 | 0.260 | 14.119 | 0.176 | na | 0 |
| **MSTRG.18255\|evm.TU.Herato1803.22\|evm.TU.Herato1803.28** | 8.531 | 6.121 | 28.630 | 0.017 | 1.914 | 6.453 | 35.671 | 0.119 | circadian clock-controlled protein-like [Vanessa tameamea] | 2.00E-64 |
| **MSTRG.17035\|evm.TU.Herato1701.207_evm.TU.Herato1701.208_evm.TU.Herato1701.210** | 1.750 | 4.257 | 27.600 | 0.017 | 0.705 | 4.558 | 17.744 | 0.149 | TPA_inf: cytochrome P450 CYP405A4 [Heliconius erato] | 0 |
| **MSTRG.21292\|evm.TU.Herato2001.642** | 2.802 | 5.208 | 26.958 | 0.018 | 0.651 | 5.776 | 26.047 | 0.090 | serine protease snake-like [Vanessa tameamea] | 2.00E-89 |
| **MSTRG.18256*** | **7.243** | **1.525** | **24.987** | **0.021** | **2.406** | **1.762** | **65.210** | **0.024** | **na** | **na** |
| **MSTRG.20971\|evm.TU.Herato2001.420** | 2.234 | 6.824 | 21.519 | 0.027 | 0.420 | 7.324 | 17.834 | 0.144 | fringe [Junonia coenia] | 0.00E+00 |
| **MSTRG.9120\|evm.TU.Herato1003.15_evm.TU.Herato1003.17\|evm.TU.Herato1003.22\|evm.TU.Herato1003.23** | 1.194 | 4.395 | 19.771 | 0.031 | 0.396 | 4.704 | 14.415 | 0.175 | protein MLP1 homolog isoform X2 [Aphantopus hyperantus] | 0 |
| **MSTRG.3894\|evm.TU.Herato0503.238** | **0.944** | **4.236** | **19.284** | **0.032** | **0.738** | **4.322** | **44.080** | **0.044** | **chitin deacetylase 1 [Zerene cesonia]** | **0.00E+00** |

| | | | | | | | | | | |
|---|---|---|---|---|---|---|---|---|---|---|
| **MSTRG.681\|evm.TU.Herato0101.519** | 0.788 | 4.675 | 18.055 | 0.037 | 0.397 | 4.839 | 18.989 | 0.131 | chromatin complexes subunit BAP18-like [Aphantopus hyperantus] | 2.00E-48 |
| **MSTRG.10736\|novel_gene_126** | 4.930 | 1.026 | 16.283 | 0.045 | 1.236 | 1.592 | 12.718 | 0.190 | Transposon Ty3-G Gag-Pol polyprotein [Operophtera brumata] | 1.00E-170 |
| **MSTRG.2116\|evm.TU.Herato0215.126** | 4.241 | 4.681 | 16.239 | 0.045 | 0.926 | 5.107 | 19.894 | 0.149 | calcium/calmodulin-dependent protein kinase kinase 1 [Aphantopus hyperantus] | 0.00E+00 |
| **MSTRG.2111\|evm.TU.Herato0215.124** | 4.881 | 2.985 | 14.978 | 0.052 | 1.103 | 3.502 | 20.259 | 0.131 | uncharacterized protein LOC113404666 [Vanessa tameamea] | 7.00E-32 |
| **MSTRG.17608\|evm.TU.Herato1706.3** | 1.721 | 3.190 | 14.614 | 0.053 | 1.477 | 3.291 | 90.382 | 0.011 | PREDICTED: tetraspanin-9-like [Papilio polytes] | 4.00E-70 |
| **MSTRG.17430\|evm.TU.Herato1704.13** | 1.546 | 4.868 | 14.176 | 0.057 | 0.569 | 5.158 | 29.804 | 0.083 | neprilysin-4-like [Aphantopus hyperantus] | 0.00E+00 |
| **MSTRG.6129\|evm.TU.Herato0701.205** | 2.086 | 2.768 | 14.074 | 0.057 | 0.683 | 3.212 | 12.873 | 0.189 | solute carrier family 28 member 3-like [Bicyclus anynana] | 0.00E+00 |

| | | | | | | | | | | |
|---|---|---|---|---|---|---|---|---|---|---|
| **MSTRG.1644\|evm.TU.Herato0211.46** | 1.076 | 4.242 | 13.616 | 0.059 | 0.589 | 4.413 | 37.831 | 0.055 | cell cycle control protein 50A-like isoform X2 [Vanessa tameamea] | 5.00E-78 |
| **MSTRG.14815\|evm.TU.Herato1411.237** | 0.648 | 4.162 | 12.524 | 0.069 | 0.413 | 4.258 | 14.582 | 0.175 | pyroglutamyl-peptidase 1 [Vanessa tameamea] | 7.00E-131 |
| **MSTRG.7867\|evm.TU.Herato0821.114** | 2.116 | 5.488 | 12.238 | 0.073 | 0.348 | 6.047 | 13.712 | 0.179 | putative phospholipase B-like 2 [Vanessa tameamea] | 0.00E+00 |
| **MSTRG.12474** | 5.134 | 1.628 | 12.111 | 0.074 | 0.683 | 2.338 | 12.258 | 0.196 | na | na |
| **MSTRG.12267\|evm.TU.Herato1202.501** | 1.625 | 4.813 | 11.141 | 0.083 | 0.483 | 5.209 | 21.059 | 0.117 | nanos protein [Danaus plexippus plexippus] | 1.00E-144 |
| **MSTRG.12213\|evm.TU.Herato1202.469** | 0.824 | 7.357 | 10.872 | 0.085 | 0.425 | 7.528 | 16.759 | 0.151 | Putative polypeptide N-acetylgalactosaminyl transferase 9 [Papilio machaon] | 0.00E+00 |
| **MSTRG.21139\|evm.TU.Herato2001.552** | 0.543 | 5.284 | 10.651 | 0.088 | 0.289 | 5.398 | 13.075 | 0.187 | selenoprotein M-like [Danaus plexippus plexippus] | 2.00E-46 |
| **MSTRG.4596** | 1.523 | -0.825 | 9.588 | 0.103 | 4.611 | -1.120 | 23.398 | 0.099 | na | na |

| | | | | | | | | | | |
|---|---|---|---|---|---|---|---|---|---|---|
| **MSTRG.20663\|evm.TU.Herato2001.192** | 0.771 | 2.995 | 9.229 | 0.108 | 0.506 | 3.109 | 14.529 | 0.175 | BTB/POZ domain-containing adapter for CUL3-mediated RhoA degradation protein 3 [Trichoplusia ni] | 0.00E+00 |
| **MSTRG.21393\|evm.TU.Herato2001.698\|evm.TU.Herato2001.699** | 0.851 | 3.974 | 7.964 | 0.132 | 0.469 | 4.158 | 14.275 | 0.175 | acetyl-CoA carboxylase isoform X1 [Vanessa tameamea] | 0.00E+00 |
| **MSTRG.9217\|evm.TU.Herato1003.75** | 1.841 | 0.925 | 7.907 | 0.133 | 0.993 | 1.159 | 13.247 | 0.184 | hydroxylysine kinase [Vanessa tameamea] | 0.00E+00 |
| **MSTRG.875\|evm.TU.Herato0101.675\|evm.TU.Herato0101.677** | 0.516 | 5.310 | 7.848 | 0.134 | 0.328 | 5.393 | 14.497 | 0.175 | cysteine protease ATG4B [Bicyclus anynana] | 0.00E+00 |
| **MSTRG.7970\|evm.TU.Herato0901.44** | 2.371 | 2.202 | 7.773 | 0.137 | 0.647 | 2.721 | 13.922 | 0.176 | IDLSRF-like peptide [Bicyclus anynana] | 2.00E-131 |
| **MSTRG.641\|evm.TU.Herato0101.485** | 0.648 | 9.456 | 7.084 | 0.152 | 0.346 | 9.578 | 12.555 | 0.192 | dopa decarboxylase [Heliconius melpomene malleti] | 0.00E+00 |
| **MSTRG.20662\|evm.TU.Herato2001.191** | 0.462 | 4.393 | 6.902 | 0.157 | 0.695 | 4.305 | 54.035 | 0.032 | INO80 complex subunit B isoform X1 [Ostrinia furnacalis] | 7.00E-168 |

| | | | | | | | | | | |
|---|---|---|---|---|---|---|---|---|---|---|
| **MSTRG.18061\|evm.TU.Herato1801.88** | 0.429 | 5.314 | 6.878 | 0.157 | 0.674 | 5.217 | 44.191 | 0.044 | protein TANC2 isoform X2 [Vanessa tameamea] | 0.00E+00 |
| **MSTRG.16897\|evm.TU.Herato1701.114** | 0.557 | 8.223 | 6.714 | 0.161 | 0.650 | 8.200 | 21.925 | 0.111 | flocculation protein FLO11 isoform X1 [Danaus plexippus plexippus] | 0.00E+00 |
| **MSTRG.19877\|evm.TU.Herato1908.46** | 1.716 | 4.514 | 6.406 | 0.172 | 0.821 | 4.810 | 17.792 | 0.155 | uncharacterized protein LOC113404717 [Vanessa tameamea] | 0.00E+00 |
| **MSTRG.17497\|evm.TU.Herato1705.21** | 2.474 | 6.833 | 6.075 | 0.182 | 0.643 | 7.310 | 40.304 | 0.050 | uncharacterized protein LOC112045811 [Bicyclus anynana] | 2.00E-51 |
| **MSTRG.20088\|evm.TU.Herato1910.89\|evm.TU.Herato1910.91** | -1.177 | 4.707 | 43.197 | 0.007 | -0.457 | 4.258 | 15.312 | 0.165 | transmembrane protein 136-like [Vanessa tameamea] | 8.00E-128 |
| **MSTRG.21343\|evm.TU.Herato2001.670_evm.TU.Herato2001.671** | -1.774 | 4.151 | 38.396 | 0.009 | -1.032 | 3.623 | 25.269 | 0.098 | glucose-1-phosphatase-like [Aphantopus hyperantus] | 7.00E-153 |
| **MSTRG.14548\|evm.TU.Herato1411.67** | -1.259 | 3.280 | 31.228 | 0.014 | -0.936 | 3.047 | 25.298 | 0.092 | hepatocyte nuclear factor 4-gamma isoform X3 [Aphantopus hyperantus] | 0.00E+00 |

| | | | | | | | | | | |
|---|---|---|---|---|---|---|---|---|---|---|
| **MSTRG.5330\|evm.TU.Herato0606.247** | -2.400 | 4.490 | 28.716 | 0.017 | -1.710 | 4.114 | 37.997 | 0.086 | calcium-activated potassium channel slowpoke isoform X31 [Vanessa tameamea] | 0.00E+00 |
| **MSTRG.10838\|evm.TU.Herato1108.274\|evm.TU.Herato1108.275\|evm.TU.Herato1108.277** | -0.788 | 6.332 | 25.518 | 0.019 | -0.524 | 6.182 | 33.820 | 0.068 | Multidrug resistance-associated protein 4 [Papilio machaon] | 1.00E-35 |
| **MSTRG.5161\|evm.TU.Herato0310.312\|evm.TU.Herato0310.313** | -0.891 | 4.599 | 17.517 | 0.039 | -0.437 | 4.327 | 16.858 | 0.151 | no hits | na |
| **MSTRG.12481\|evm.TU.Herato1202.627** | -2.595 | 0.616 | 13.765 | 0.059 | -1.489 | -0.295 | 13.382 | 0.182 | condensin complex subunit 1 isoform X1 [Vanessa tameamea] | 0.00E+00 |
| **MSTRG.13621\|evm.TU.Herato1301.564** | -0.513 | 6.695 | 12.199 | 0.072 | -0.767 | 6.855 | 45.451 | 0.043 | Krueppel homolog 2 [Vanessa tameamea] | 2.00E-162 |
| **MSTRG.1813\|evm.TU.Herato0213.2** | -1.752 | 5.625 | 12.050 | 0.075 | -0.352 | 4.667 | 13.895 | 0.176 | luciferase-like 8, partial [Heliconius erato] | 0.00E+00 |
| **MSTRG.15493\|evm.TU.Herato1507.212** | -0.974 | 4.880 | 9.197 | 0.109 | -0.374 | 4.510 | 13.676 | 0.179 | hypothetical protein evm_013227 [Chilo suppressalis] | 8.00E-70 |
| **MSTRG.4075\|evm.TU.Herato0508.49** | -0.811 | 5.537 | 8.785 | 0.115 | -0.520 | 5.385 | 32.559 | 0.073 | hypothetical protein RR46_04746 [Papilio xuthus] | 0.00E+00 |

| | | | | | | | | | | |
|---|---|---|---|---|---|---|---|---|---|---|
| **MSTRG.604\|evm.TU.Herato0101.456** | -0.714 | 5.993 | 8.591 | 0.119 | -0.604 | 5.935 | 15.093 | 0.175 | uncharacterized protein LOC112050048 [Bicyclus anynana] | 0.00E+00 |
| **MSTRG.11531\|evm.TU.Herato1202.83\|evm.TU.Herato1202.84** | -0.796 | 3.092 | 6.519 | 0.167 | -0.529 | 2.939 | 12.675 | 0.190 | PREDICTED: histone H2B [Plutella xylostella] | 6.00E-82 |
| **MSTRG.12897\|evm.TU.Herato1301.53** | -0.712 | 5.143 | 6.253 | 0.175 | -0.553 | 5.059 | 18.032 | 0.143 | protein espinas isoform X3 [Vanessa tameamea] | 0.00E+00 |
| **MSTRG.14849\|evm.TU.Herato1501.1** | -2.660 | 4.687 | 6.024 | 0.184 | -0.839 | 3.238 | 28.371 | 0.086 | chaoptin-like [Danaus plexippus plexippus] | 0.00E+00 |
| **MSTRG.6831\|evm.TU.Herato0701.670** | -2.363 | 1.531 | 5.793 | 0.192 | -1.505 | 0.654 | 12.125 | 0.198 | cysteine and histidine-rich protein 1-like [Vanessa tameamea] | 0.00E+00 |
| ***Heliconius melpomene*** **DAY 5** | | | | | | | | | | |
| **MSTRG.18872\|HMEL003022g1** | 1.494 | 2.443 | 17.566 | 0.107 | 0.877 | 2.619 | 52.117 | 0.158 | LOW QUALITY PROTEIN: xanthine dehydrogenase 1-like [Bicyclus anynana] | 0 |
| **DAY 7** | | | | | | | | | | |

| | | | | | | | | | | |
|---|---|---|---|---|---|---|---|---|---|---|
| **MSTRG.14267\|HMEL031592g1** | 1.747 | 2.525 | 46.347 | 0.057 | 0.819 | 2.749 | 18.494 | 0.195 | uncharacterized protein LOC113397890 [Vanessa tameamea] | 4.00E-87 |
| **MSTRG.8396\|HMEL013392g1\|HMEL037724g1** | 0.815 | 3.116 | 16.497 | 0.146 | 0.544 | 3.218 | 21.269 | 0.151 | drebrin-like protein B [Vanessa tameamea] | 1.00E-177 |
| **MSTRG.8785\|HMEL002509g1** | 0.808 | 3.784 | 15.115 | 0.160 | 0.657 | 3.840 | 23.923 | 0.131 | hemicentin-1-like isoform X1 [Vanessa tameamea] | 0 |
| **MSTRG.2699\|HMEL002124g1** | 1.398 | 6.822 | 14.991 | 0.161 | 0.718 | 7.016 | 21.448 | 0.173 | probable fatty acid-binding protein [Danaus plexippus plexippus] | 9.00E-66 |
| **MSTRG.4246\|HMEL036361g1** | 1.578 | 2.876 | 14.618 | 0.166 | 1.219 | 2.937 | 27.330 | 0.120 | cuticle protein 8-like [Bicyclus anynana] | 2.00E-72 |
| **MSTRG.18497\|HMEL012022g1** | 2.982 | 1.492 | 14.528 | 0.167 | 4.471 | 1.288 | 106.926 | 0.004 | brachyurin-like [Aphantopus hyperantus] | 1.00E-109 |
| **MSTRG.18976\|HMEL008071g1\|HMEL008071g2** | 1.182 | 1.740 | 14.134 | 0.170 | 1.494 | 1.656 | 35.183 | 0.054 | tetra-peptide repeat homeobox protein 1-like [Vanessa tameamea] | 4.00E-37 |
| **MSTRG.20941\|HMEL034284g1** | -4.999 | 2.881 | 21.651 | 0.113 | -2.481 | 0.775 | 23.140 | 0.143 | circadian clock-controlled protein-like [Vanessa tameamea] | 5.00E-27 |

Table S14: Concordantly expressed genes between *H. erato* and *H. melpomene* with an FDR < 0.2. Bold indicates genes which are significantly DE expressed (FDR < 0.05) in both species.

| erato.gene | logFC.erato | logCPM.erato | FDR.erato | mel.gene | identity(%) | expect | logFC.mel | logCPM | FDR.mel | NCBI Blastp Annotation | E-value |
|---|---|---|---|---|---|---|---|---|---|---|---|
| **Race** | | | | | | | | | | | |
| **Day 5** | | | | | | | | | | | |
| evm.model.Herato0606.92 | 2.312 | 1.730 | 0.061 | HMEL016763g1 | 95.639 | 0 | 1.099 | 3.282 | 0.066 | putative carbonic anhydrase 3 [Aphantopus hyperantus] | 0 |
| evm.model.Herato2001.729 | 0.783 | 4.304 | 0.087 | HMEL004468g1 | 96.468 | 0 | 0.830 | 4.670 | 0.111 | glucose-6-phosphate exchanger SLC37A2 isoform X1 [Vanessa tameamea] | 0 |
| evm.model.Herato1805.232 | 0.650 | 5.655 | 0.119 | HMEL006054g1 | 96.678 | 0 | 0.634 | 5.469 | 0.081 | solute carrier family 12 member 9 isoform X2 [Aphantopus hyperantus] | 0 |
| evm.model.Herato2001.75 | 0.618 | 5.295 | 0.125 | HMEL017055g1 | 97.297 | 7.08E-134 | 1.083 | 4.782 | 0.037 | peroxisomal membrane protein 2 [Bicyclus anynana] | 3.00E-122 |
| evm.model.Herato1901.116_evm.model.Herato1901.120 | 1.085 | 3.358 | 0.147 | HMEL010934 | 94.375 | 0 | 1.617 | 3.283 | 0.037 | cytochrome P450 4C1-like [Danaus plexippus plexippus] | 9.00E-157 |
| evm.model.Herato0503.162 | 1.450 | 2.221 | 0.150 | HMEL008925g1 | 93.986 | 0 | 2.697 | 0.757 | 0.097 | pickpocket protein 28 [Vanessa tameamea] | 0 |
| evm.model.Herato0701.776 | -7.725 | 1.805 | 0.003 | HMEL037763g1 | 86.275 | 2.12E-08 | -1.282 | 1.421 | 0.075 | reticulon-4-interacting protein 1, mitochondrial [Bombyx mori] | 1.00E-32 |
| evm.model.Herato1805.48 | -0.884 | 3.619 | 0.011 | HMEL003070g1 | 97.035 | 0 | -0.544 | 3.552 | 0.162 | threonine aspartase 1 [Vanessa tameamea] | 0 |
| evm.model.Herato0801.233 | -1.429 | 2.805 | 0.017 | HMEL038031g1 | 92.492 | 0 | -1.067 | 2.867 | 0.072 | mitochondrial amidoxime-reducing component 1 isoform X2 [Manduca sexta] | 0 |

| | | | | | | | | | | | |
|---|---|---|---|---|---|---|---|---|---|---|---|
| **evm.model.Herato1004.1** | **-0.977** | **3.717** | **0.026** | **HMEL030760g1** | **95.021** | **1.26E-174** | **-2.058** | **2.919** | **0.037** | **Gamma-glutamyl cyclotransferase-like venom protein isoform 1, partial [Operophtera brumata]** | **4.00E-108** |
| **evm.model.Herato0701.772** | **-1.454** | **2.037** | **0.029** | **HMEL005305g3** | **94.811** | **0** | **-1.613** | **1.856** | **0.035** | **Fatty acid synthase [Danaus plexippus plexippus]** | **0** |
| evm.model.Herato1901.44 | -2.216 | 3.225 | 0.058 | HMEL034227g1 | 62.048 | 0 | -2.664 | 1.676 | 0.129 | PREDICTED: uncharacterized protein LOC106710892 [Papilio machaon] | 0 |
| evm.model.Herato1301.53 | -0.928 | 5.189 | 0.084 | HMEL031781g1 | 99.315 | 0 | -0.601 | 4.465 | 0.172 | protein espinas isoform X3 [Vanessa tameamea] | 0 |
| evm.model.Herato1005.109 | -0.646 | 3.735 | 0.142 | HMEL013306g1 | 83.571 | 1.07E-76 | -0.735 | 3.875 | 0.135 | PREDICTED: E3 ubiquitin-protein ligase RNF4-like [Papilio polytes] | 2.00E-39 |
| evm.model.Herato1005.218 | -0.791 | 4.931 | 0.145 | HMEL030866g1 | 96.407 | 0 | -1.158 | 2.614 | 0.120 | myrosinase 1-like [Vanessa tameamea] | 0 |
| evm.model.Herato1301.541 | -0.719 | 3.126 | 0.164 | HMEL009927g1 | 68.487 | 0 | -1.047 | 3.384 | 0.041 | putative pre-mRNA-splicing factor ATP-dependent RNA helicase DHX16 [Vanessa tameamea] | 1.00E-64 |
| evm.model.Herato0606.27 | -0.746 | 2.997 | 0.192 | HMEL008899g1 | 95.299 | 4.87E-168 | -2.198 | 3.004 | 0.040 | leucine-rich melanocyte differentiation-associated protein-like [Vanessa tameamea] | 5.00E-136 |
| **Day 7** | | | | | | | | | | | |
| evm.model.Herato0701.610 | 1.238 | 1.873 | 0.153 | HMEL037635g1 | 92.13 | 0 | 1.084 | 3.409 | 0.066 | sperm-associated antigen 6-like [Ostrinia furnacalis] | 0 |
| evm.model.Herato1004.1 | -1.675 | 4.087 | 0.004 | HMEL030760g1 | 95.021 | 0 | -1.333 | 2.991 | 0.060 | Gamma-glutamyl cyclotransferase-like venom protein isoform 1, partial [Operophtera brumata] | 0 |
| evm.model.Herato0701.716 | -1.399 | 2.279 | 0.019 | HMEL037725g1 | 65.987 | 0 | -1.276 | 2.659 | 0.172 | protein ZDS1-like [Vanessa tameamea] | 0 |
| evm.model.Herato0204.6 | -2.914 | 1.107 | 0.019 | HMEL022589g1 | 97.268 | 0 | -3.063 | 1.582 | 0.060 | TWiK family of potassium channels protein 12 [Aphantopus hyperantus] | 0 |
| evm.model.Herato0419.55 | -2.078 | 5.269 | 0.022 | HMEL013922g1 | 70.292 | 0 | -0.523 | 3.641 | 0.189 | ATP-dependent Clp protease ATP-binding subunit clpX-like, mitochondrial isoform X1 [Vanessa tameamea] | 0 |
| evm.model.Herato0214.16 | -4.442 | 4.578 | 0.026 | HMEL014358g1 | 81.265 | 0 | -0.615 | 6.778 | 0.113 | E3 ubiquitin-protein ligase SIAH1-like isoform X3 [Bicyclus anynana] | 0 |

| | | | | | | | | | | | |
|---|---|---|---|---|---|---|---|---|---|---|---|
| evm.model.Herato1108.426 | -0.713 | 7.556 | 0.078 | HMEL009792g1 | 98.282 | 0 | -0.575 | 7.463 | 0.117 | dynamin isoform X10 [Aphantopus hyperantus] | 0 |
| evm.model.Herato0801.322 | -0.954 | 4.645 | 0.091 | HMEL010036g1 | 98.131 | 0 | -0.770 | 2.715 | 0.152 | 28S ribosomal protein S5, mitochondrial [Aphantopus hyperantus] | 0 |
| evm.model.Herato2101.87 | -0.434 | 6.700 | 0.133 | HMEL013502g1 | 98.688 | 0 | -1.433 | 6.768 | 0.090 | zinc-type alcohol dehydrogenase-like protein C1773.06c [Manduca sexta] | 0 |
| evm.model.Herato0205.2 | -3.573 | 2.130 | 0.147 | HMEL033207g1 | 80.782 | 0 | -2.253 | 2.086 | 0.082 | uncharacterized protein LOC112049287 [Bicyclus anynana] | 0 |
| evm.model.Herato1805.220 | -1.434 | 3.752 | 0.149 | HMEL003246g1 | 95.722 | 0 | -1.578 | 1.672 | 0.198 | organic cation transporter protein-like [Vanessa tameamea] | 0 |
| evm.model.Herato0211.81 | -0.585 | 3.613 | 0.154 | HMEL015454g1 | 95.873 | 0 | -0.766 | 2.478 | 0.170 | transcription initiation protein SPT3 homolog [Vanessa tameamea] | 0 |
| evm.model.Herato1005.61 | -1.642 | 0.595 | 0.167 | HMEL005932 | 95.519 | 0 | -2.696 | 2.571 | 0.063 | cytochrome P450 4C1-like [Aphantopus hyperantus] | 0 |
| **Tissue** | | | | | | | | | | | |
| **Day 5** | | | | | | | | | | | |
| evm.model.Herato1001.152 | 3.334 | 0.505 | 0.149 | HMEL002901 | 99.296 | 0 | 2.338 | 1.044 | 0.041 | paired box pox-neuro protein [Vanessa tameamea] | 0 |
| evm.model.Herato0801.87 | -1.839 | 3.853 | 0.080 | HMEL037907g1 | 100 | 0 | -1.807 | 3.979 | 0.044 | protein doublesex-like isoform X1 [Vanessa tameamea] | 0 |
| **Day 7** | | | | | | | | | | | |
| evm.model.Herato0701.190 | 1.949 | 11.511 | 0.032 | HMEL007136g1 | 93.377 | 0 | 1.404 | 9.371 | 0.163 | cuticle protein 1-like [Danaus plexippus plexippus] | 0 |
| **evm.model.Herato0701.220** | **1.906** | **2.423** | **0.034** | **HMEL009343g2** | **98.638** | **0** | **1.702** | **2.967** | **0.023** | **homeobox protein invected-like isoform X1 [Vanessa tameamea]** | **0** |
| evm.model.Herato2101.383 | 1.192 | 2.494 | 0.060 | HMEL008471g6 | 99.407 | 0 | 1.575 | 2.236 | 0.032 | homeobox protein araucan-like isoform X2 [Vanessa tameamea] | 0 |
| evm.model.Herato1003.176 | 0.573 | 7.077 | 0.074 | HMEL030728g1 | 99.643 | 0 | 0.925 | 7.318 | 0.003 | protein limb expression 1 homolog isoform X1 [Vanessa tameamea] | 0 |
| evm.model.Herato0208.10 | 1.296 | 1.458 | 0.075 | HMEL017792g5 | 94.187 | 0 | 0.616 | 3.141 | 0.168 | uncharacterized protein LOC117985176 [Aphantopus hyperantus] | 0 |

| | | | | | | | | | | | |
|---|---|---|---|---|---|---|---|---|---|---|---|
| evm.model.Herato0901.76 | 1.364 | 2.272 | 0.082 | HMEL008230g1 | 99.552 | 0 | 1.159 | 1.348 | 0.163 | uncharacterized protein LOC113401684 isoform X1 [Vanessa tameamea] | 0 |
| evm.model.Herato1704.13 | 0.569 | 5.158 | 0.083 | HMEL017687g1 | 80.077 | 0 | 0.583 | 5.852 | 0.053 | neprilysin-4-like [Aphantopus hyperantus] | 0 |
| evm.model.Herato0211.105 | 0.442 | 6.843 | 0.088 | HMEL002124g1 | 93.617 | 0 | 0.718 | 7.016 | 0.173 | probable fatty acid-binding protein [Danaus plexippus plexippus] | 0 |
| evm.model.Herato0701.563 | 0.718 | 7.387 | 0.093 | HMEL021853g1 | 98.413 | 0 | 0.702 | 6.947 | 0.020 | cuticle protein 18.6-like [Bicyclus anynana] | 0 |
| evm.model.Herato1301.179 | 0.984 | 2.057 | 0.097 | HMEL007991g4 | 99.471 | 0 | 0.865 | 3.134 | 0.166 | cuticle protein 7-like [Vanessa tameamea] | 0 |
| evm.model.Herato1301.115 | 0.846 | 2.279 | 0.108 | HMEL015354g1 | 99.568 | 0 | 0.838 | 1.964 | 0.186 | GATA-binding factor C-like isoform X2 [Vanessa tameamea] | 0 |
| evm.model.Herato0101.722 | 0.425 | 6.064 | 0.128 | HMEL011421g3 | 92.096 | 0 | 1.539 | 3.113 | 0.007 | tubulin glycylase 3A-like isoform X1 [Vanessa tameamea] | 0 |
| evm.model.Herato0901.205 | 0.462 | 9.382 | 0.131 | HMEL014632g1 | 95.028 | 0 | 0.889 | 9.432 | 0.012 | endochitinase [Vanessa tameamea] | 0 |
| evm.model.Herato1001.42 | 0.536 | 9.577 | 0.132 | HMEL017377g1 | 96.727 | 0 | 0.431 | 8.496 | 0.139 | chymotrypsin-1-like [Vanessa tameamea] | 0 |
| evm.model.Herato1301.86 | -1.127 | 3.618 | 0.020 | HMEL031805g1 | 98.2 | 0 | -0.467 | 5.026 | 0.121 | facilitated trehalose transporter Tret1-like isoform X1 [Vanessa tameamea] | 0 |
| **evm.model.Herato1411.170** | **-0.738** | **8.444** | **0.027** | **HMEL005529g1** | **98.582** | **0** | **-0.762** | **9.122** | **0.005** | **hemocyte protein-glutamine gamma-glutamyltransferase-like [Vanessa tameamea]** | **0** |
| **evm.model.Herato1007.62** | **-1.216** | **7.046** | **0.027** | **HMEL015975g1** | **93.103** | **0** | **-0.795** | **3.002** | **0.047** | **uncharacterized protein LOC117993154 [Aphantopus hyperantus]** | **0** |
| evm.model.Herato0101.17 | -0.740 | 9.229 | 0.037 | HMEL022617g1 | 94.937 | 0 | -0.444 | 9.207 | 0.112 | GATA zinc finger domain-containing protein 14-like [Aphantopus hyperantus] | 0 |
| **evm.model.Herato0801.87** | **-0.884** | **3.032** | **0.049** | **HMEL037907g1** | **100** | **0** | **-1.676** | **4.105** | **0.006** | **protein doublesex-like isoform X1 [Vanessa tameamea]** | **0** |

| | | | | | | | | | | | |
|---|---|---|---|---|---|---|---|---|---|---|---|
| evm.model.Herato0601.10 | -0.536 | 7.260 | 0.084 | HMEL012144g1 | 99.099 | 0 | -0.428 | 6.486 | 0.151 | zinc finger protein Noc [Vanessa tameamea] | 0 |
| evm.model.Herato2101.123 | -0.633 | 5.067 | 0.144 | HMEL008807g1 | 84.559 | 0 | -0.566 | 5.264 | 0.168 | tan [Heliconius melpomene malleti] | 0 |
| evm.model.Herato1301.434 | -0.434 | 8.426 | 0.151 | HMEL008086g1 | 98.131 | 0 | -0.468 | 9.012 | 0.142 | cuticle protein 19.8 isoform X1 [Vanessa tameamea] | 0 |

Table S15. Genes differentially expressed (FDR<0.05) between *H. erato cyrbia* and *H. erato demophoon* within the QTL intervals.

| Phenotype | Stage | gene id | annotated gene | logFC | logCPM | FDR | scaffold | start | stop | protein hit (FlyBase) | comment |
|---|---|---|---|---|---|---|---|---|---|---|---|
| **Chr 20** | | | | | | | | | | | |
| **BR** | **Day 5** | MSTRG.21120 | | -1.365 | 3.047 | 0.034 | Herato2001 | 13665388 | 13667572 | tho2 | immediately before (392bp) tho2 (Herato2001: 13,667,964-13,668,985) , poor hit (E value > 1) |
| **BR** | | MSTRG.21053 | evm.TU.Herato2001.480 | -0.738 | 6.529 | 0.035 | Herato2001 | 12606947 | 12613794 | Acyl-CoA binding protein 1 | na |
| **BR** | **Day 7** | MSTRG.21108 | evm.TU.Herato2001.529 | 1.914 | 4.091 | 0.005 | Herato2001 | 13118144 | 13142065 | Dmel\CG11318 | na |
| **BR** | | MSTRG.20935 | | 8.806 | -0.216 | 0.006 | Herato2001 | 9817864 | 9818213 | Dmel\CG10904 | large distance from genomic feature, downstream of Dmel\CG10904 (Herato2001: 9,835,885-9,839,014) |
| **BR** | | MSTRG.21007 | | 3.730 | 1.757 | 0.011 | Herato2001 | 11641700 | 11643873 | no hit | exact match |
| **BR** | | MSTRG.21045 | evm.TU.Herato2001.474 | -8.527 | -0.402 | 0.022 | Herato2001 | 12513101 | 12535184 | Tetraspanin 26A | na |
| **BR** | | MSTRG.20971 | evm.TU.Herato2001.420 | 2.234 | 6.824 | 0.027 | Herato2001 | 10786686 | 10804234 | fringe | na |
| **Chr Z** | | | | | | | | | | | |
| **BR, LUM, RS** | **Day 5** | MSTRG.21995 | | -9.396 | 0.264 | 0.003 | Herato2101 | 10209394 | 10209747 | ATPase 8B | nearest gene 709 bp upstream |
| **BR, LUM, RS** | | MSTRG.22090 | evm.TU.Herato2101.409 | -9.324 | 0.173 | 0.004 | Herato2101 | 12435747 | 12453213 | super sex combs | na |
| **BR, LUM, RS** | | MSTRG.22036 | | -1.129 | 4.704 | 0.006 | Herato2101 | 10864412 | 10867976 | wacky | wacky (Herato2101: 10,868,052-10,881,937 ) ; Dynein heavy chain at 16F (Herato2101: 10,853,504-10,864,142) |
| **BR, LUM, RS** | | MSTRG.21947 | | -2.230 | 2.952 | 0.008 | Herato2101 | 9228419 | 9229979 | dunce | downstream dunce Herato2101: 9,232,569-9,323,532 ) |

| | | | | | | | | | | |
|---|---|---|---|---|---|---|---|---|---|---|
| **BR, LUM, RS** | MSTRG.2 1922 | evm.TU.Herato2101.296 | 1.846 | 2.644 | 0.010 | Herato2101 | 8312400 | 8313137 | no hit | na |
| **RS** | MSTRG.2 1759 | evm.TU.Herato2101.179 | -1.896 | 3.495 | 0.011 | Herato2101 | 5162891 | 5181891 | Polypeptide N-acetylgalactosaminyl transferase 35A | na |
| **BR, LUM, RS** | MSTRG.2 1925 | | -2.405 | 1.881 | 0.010 | Herato2101 | 8391373 | 8393792 | Dmel\CG7888 | exact match |
| **BR, LUM, RS** | MSTRG.2 1910 | | -1.088 | 3.331 | 0.013 | Herato2101 | 8128295 | 8129254 | moleskin | upstream of moleskin (Herato2101: 8,116,509-8,124,979 ) |
| **BR, LUM, RS** | MSTRG.2 2060 | | 1.523 | 3.241 | 0.015 | Herato2101 | 11644379 | 11647064 | Dmel\CG18659 | upstream of Herato2101: 11,595,296-11,638,782 |
| **BR, LUM** | MSTRG.2 2097 | | -4.799 | -1.420 | 0.015 | Herato2101 | 12536727 | 12612327 | Glutamate receptor IA | very large transcript, end overlaps with Glutamate receptor IA( Herato2101: 12,610,251-12,656,299) |
| **BR, LUM, RS** | MSTRG.2 1992 | | -2.784 | 0.752 | 0.016 | Herato2101 | 10124805 | 10127463 | IGF-II mRNA-binding protein | large distance from genomic feature, closest to IGF-II mRNA-binding protein (Herato2101: 10,133,734-10,191,778 ) |
| **BR, LUM, RS** | MSTRG.2 1899 | | -3.067 | 0.732 | 0.028 | Herato2101 | 7873746 | 7874214 | Dmel\CG42269 | large distance from genomic feature, closest to Dmel\CG42269 (Herato2101: 7,882,762-7,884,480) |
| **BR, LUM, RS** | MSTRG.2 1877 | evm.TU.Herato2101.263 | -0.728 | 3.659 | 0.029 | Herato2101 | 7365009 | 7386279 | kon-tiki | na |
| **BR, LUM, RS** | MSTRG.2 2022 | | 1.106 | 2.683 | 0.031 | Herato2101 | 10495531 | 10524827 | CG3739-PB | large transcript, includes CG3739-PB but E-value > 0.90 |
| **BR, LUM, RS** | MSTRG.2 1959 | | 2.954 | -0.441 | 0.032 | Herato2101 | 9404792 | 9405280 | Dmel\CG13293 | overlap with two genes, chose largest gene for blast (Dmel\CG13293 Herato2101: 9,339,580-9,409,387 ) |
| **BR, LUM, RS** | MSTRG.2 1897 | | -2.592 | -0.391 | 0.033 | Herato2101 | 7853355 | 7853655 | Transient receptor potential cation channel A1 | exact match |

| | | | | | | | | | | | |
|---|---|---|---|---|---|---|---|---|---|---|---|
| **RS** | | MSTRG.21743 | | -1.217 | 2.834 | 0.037 | Herato2101 | 4973958 | 4975233 | Site-1 protease | 291 bp downstream of Site-1 protease (Herato2101: 4,975,524-4,997,675) |
| **BR, LUM, RS** | | MSTRG.22035 | evm.TU.Herato2101.375 | 3.459 | 0.192 | 0.043 | Herato2101 | 10815026 | 10848055 | Dynein heavy chain at 16F | na |
| **BR, LUM, RS** | | MSTRG.21867 | evm.TU.Herato2101.255 | 1.465 | 2.898 | 0.044 | Herato2101 | 7114906 | 7119901 | Dmel\CG5541 | na |
| **LUM, RS** | | MSTRG.21843 | | 3.250 | 2.182 | 0.001 | Herato2101 | 6628831 | 6630351 | Dmel\CG41520 | exact match |
| **LUM, RS** | | MSTRG.21839 | | -1.741 | 3.493 | 0.006 | Herato2101 | 6605080 | 6607246 | no hit | large distance from genomic feature, downstream of Herato2101: 6,614,966-6,621,785 |
| **LUM, RS** | | MSTRG.21793 | evm.TU.Herato2101.200 | -1.137 | 5.297 | 0.007 | Herato2101 | 5793288 | 5802200 | Quiescin sulfhydryl oxidase 1 | na |
| **LUM, RS** | | MSTRG.21832 | | -1.331 | 3.298 | 0.009 | Herato2101 | 6516431 | 6525230 | Autophagy-related 9 | downstream (1491bp) Autophagy-related 9 (Herato2101: 6,526,721-6,538,967) |
| **LUM, RS** | | MSTRG.21803 | | -5.262 | 0.125 | 0.017 | Herato2101 | 5980907 | 5981143 | biniou | large distance from genomic feature, downstream to biniou (Herato2101: 5,991,068-6,012,882) |
| **LUM, RS** | | MSTRG.21851 | | -5.381 | -0.146 | 0.022 | Herato2101 | 6795954 | 6797473 | Phosphogluconate dehydrogenase | downstream (2666bp) Phosphogluconate dehydrogenase (Herato2101: 6,800,139-6,808,101) |
| **LUM, RS** | | MSTRG.21853 | evm.TU.Herato2101.242 | 3.305 | 0.251 | 0.047 | Herato2101 | 6811074 | 6814359 | CG42674 | na |
| **LUM, RS** | | MSTRG.21837 | | -1.041 | 3.300 | 0.047 | Herato2101 | 6585420 | 6585932 | Ubiquitin carboxy-terminal hydrolase L5 | upstream 5515bp of Ubiquitin carboxy-terminal hydrolase L5 (Herato2101: 6,580,417-6,584,825) |
| **BR, LUM, RS** | **Day 7** | MSTRG.21863 | evm.TU.Herato2101.252,evm.TU.Herato2101.253 | -2.286 | 7.336 | 0.002 | Herato2101 | 7007539 | 7033173 | trio | na |
| **BR, LUM, RS** | | MSTRG.21925 | | -2.255 | 1.924 | 0.003 | Herato2101 | 8391373 | 8393792 | Dmel\CG7888 | exact match |

| | | | | | | | | | | |
|---|---|---|---|---|---|---|---|---|---|---|
| **BR, LUM, RS** | MSTRG.2 1910 | | -1.260 | 3.781 | 0.006 | Herato2101 | 8128295 | 8129254 | moleskin | upstream of moleskin (Herato2101: 8,116,509-8,124,979 ) |
| **BR, LUM, RS** | MSTRG.2 1995 | | -8.767 | -0.195 | 0.008 | Herato2101 | 10209394 | 10209747 | ATPase 8B | nearest gene 709 bp upstream |
| **BR, LUM, RS** | MSTRG.2 1985 | | -1.279 | 3.797 | 0.012 | Herato2101 | 9964577 | 9969000 | Rho GTPase activating protein at 68F | immediately before (<200bp) Rho GTPase activating protein at 68F (Herato2101: 9,969,182-9,976,893) |
| **BR, LUM, RS** | MSTRG.2 2036 | | -0.817 | 4.101 | 0.016 | Herato2101 | 10864412 | 10867976 | wacky | wacky (Herato2101: 10,868,052-10,881,937 ) ; Dynein heavy chain at 16F (Herato2101: 10,853,504-10,864,142) |
| **BR, LUM, RS** | MSTRG.2 1947 | | -1.615 | 2.054 | 0.020 | Herato2101 | 9228419 | 9229979 | dunce | slightly downstream of dunce Herato2101: 9,232,569-9,323,532 ) |
| **RS** | MSTRG.2 1742 | evm.TU.Herato2101.164 | -2.354 | -0.332 | 0.021 | Herato2101 | 4946459 | 4952457 | Dmel\CG31717 | poor hit , E value > 1 |
| **BR, LUM, RS** | MSTRG.2 2001 | | 3.915 | -0.094 | 0.023 | Herato2101 | 10356441 | 10357252 | procollagen lysyl hydroxylase | in between two genes, downstream from procollagen lysyl hydroxylase (Herato2101: 10,359,604-10,392,288 ) |
| **BR, LUM, RS** | MSTRG.2 2022 | | 1.028 | 4.202 | 0.023 | Herato2101 | 10495531 | 10524827 | CG3739-PB | large transcript, includes CG3739-PB but E-value > 0.90 |
| **BR, LUM, RS** | MSTRG.2 1907 | evm.TU.Herato2101.284 | 1.540 | 0.950 | 0.039 | Herato2101 | 8100167 | 8115041 | Dmel\CG42271 | na |
| **BR, LUM, RS** | MSTRG.2 1899 | | -3.326 | 0.461 | 0.041 | Herato2101 | 7873746 | 7874214 | Dmel\CG42269 | large distance from genomic feature, closest to Dmel\CG42269 (Herato2101: 7,882,762-7,884,480) |
| **BR, LUM, RS** | MSTRG.2 2076 | evm.TU.Herato2101.401 | -4.220 | -1.176 | 0.044 | Herato2101 | 12146767 | 12154685 | Dmel\CG32260 | na |
| **BR, LUM, RS** | MSTRG.2 1964 | evm.TU.Herato2101.324 | -0.710 | 4.443 | 0.049 | Herato2101 | 9445555 | 9471870 | Dmel\CG1265 | na |
| **BR, LUM, RS** | MSTRG.2 2060 | | 2.058 | 3.282 | 0.049 | Herato2101 | 11644379 | 11647064 | Dmel\CG18659 | quite a bit upstream of Herato2101: 11,595,296-11,638,782 |

| | | | | | | | | | | |
|---|---|---|---|---|---|---|---|---|---|---|
| **LUM, RS** | MSTRG.21793 | evm.TU.Herato2101.200 | -1.392 | 6.267 | 0.001 | Herato2101 | 5793288 | 5802200 | Quiescin sulfhydryl oxidase 1 | na |
| **LUM, RS** | MSTRG.21786 | | -1.957 | 1.858 | 0.003 | Herato2101 | 5711007 | 5714276 | Kip1 ubiquitination-promoting complex subunit 1 | large distance from any genomic feature, upstream of Kip1 ubiquitination-promoting complex subunit 1 (Herato2101: 5,693,521-5,707,165) |
| **LUM, RS** | MSTRG.21787 | evm.TU.Herato2101.196 | -6.262 | -0.219 | 0.008 | Herato2101 | 5718097 | 5742135 | miles to go | poor hit , E value > 1 |
| **LUM, RS** | MSTRG.21822 | | -2.732 | 1.040 | 0.011 | Herato2101 | 6356559 | 6359488 | Dmel\CG12531 | exact match |
| **LUM, RS** | MSTRG.21843 | | 3.750 | 3.131 | 0.013 | Herato2101 | 6628831 | 6630351 | Dmel\CG41520 | exact match |
| **LUM, RS** | MSTRG.21851 | | -5.617 | 0.274 | 0.017 | Herato2101 | 6795954 | 6797473 | Phosphogluconate dehydrogenase | downstream (2666bp) Phosphogluconate dehydrogenase (Herato2101: 6,800,139-6,808,101) |
| **LUM, RS** | MSTRG.21774 | | -1.734 | 2.303 | 0.019 | Herato2101 | 5508065 | 5508627 | period | immediately downstream (72bp) of period (Herato2101: 5,508,699-5,543,290) |
| **LUM, RS** | MSTRG.21832 | | -1.935 | 5.821 | 0.022 | Herato2101 | 6516431 | 6525230 | Autophagy-related 9 | downstream (1491bp) Autophagy-related 9 (Herato2101: 6,526,721-6,538,967) |
| **LUM, RS** | MSTRG.21803 | | -2.551 | -0.259 | 0.031 | Herato2101 | 5980907 | 5981143 | biniou | large distance from genomic feature, downstream to biniou (Herato2101: 5,991,068-6,012,882) |
| **RS** | MSTRG.21751 | evm.TU.Herato2101.170 | -1.258 | 1.600 | 0.039 | Herato2101 | 5033858 | 5036106 | cricklet | na |

Table S16. Genes differentially expressed (FDR<0.2) between *H. melpomene cythera* and *H. melpomene rosina* within the QTL intervals.

| Phenotype | Stage | gene id | annotated gene | logFC | logCPM | FDR | scaffold | start | stop | protein hit (FlyBase) | novel transcript comment |
|---|---|---|---|---|---|---|---|---|---|---|---|
| **Chr 3** | | | | | | | | | | | |
| **BR** | **Day 5** | **MSTRG.3173** | | **-11.518** | **6.073** | **0.047** | **Hmel203003o** | **380684** | **383247** | **SIFamide receptor** | **downstream 653bp of SIFamide receptor (Hmel203003o: 383,900-384,613)** |
| **BR, LUM** | | MSTRG.3408 | HMEL007690g2, HMEL013806g1 | -1.825 | 2.127 | 0.131 | Hmel203003o | 3012068 | 3023317 | Heterochromatin Protein 1c | na |
| **BR. LUM** | | MSTRG.3403 | HMEL036038g1 | 1.600 | 1.836 | 0.192 | Hmel203003o | 2924821 | 3004843 | miniature | na |
| **BR** | **Day 7** | MSTRG.3297 | HMEL016174g1 | 1.168 | 3.895 | 0.060 | Hmel203003o | 1674761 | 1679707 | Dmel\CG5377 | na |
| **BR** | | MSTRG.3173 | | -11.052 | 4.856 | 0.061 | Hmel203003o | 380684 | 383247 | SIFamide receptor | downstream 653bp of SIFamide receptor (Hmel203003o: 383,900-384,613) |
| **BR** | | MSTRG.3196 | HMEL015572g1 | 1.721 | 2.951 | 0.079 | Hmel203003o | 599479 | 612509 | Saccheropin dehydrogenase 1 | na |
| **BR** | | MSTRG.3531 | HMEL036114g1 | -1.491 | 1.009 | 0.113 | Hmel203003o | 4947102 | 4949039 | pitchoune | na |
| **BR, LUM** | | MSTRG.3421 | | 6.803 | -1.539 | 0.128 | Hmel203003o | 3355012 | 3355456 | Sol1 | in region of no genomic features, equal distance between genes, usptream 8822 bp of Hmel203003o: 3,332,639-3,346,190 |
| **BR** | | MSTRG.3214 | HMEL008058g1 | 1.352 | 2.344 | 0.130 | Hmel203003o | 709863 | 710876 | Dmel\Daao1 | na |
| **BR** | | MSTRG.3151 | HMEL021694g1, HMEL021694g2 | 1.348 | 3.405 | 0.134 | Hmel203003o | 159359 | 167643 | Dmel\CG9701 | na |
| **BR** | | MSTRG.3485 | | 6.643 | -1.663 | 0.186 | Hmel203003o | 4365102 | 4365486 | Dmel\CG30413 | immediately downstream 436 bp |

| | | | | | | | | | | | |
|---|---|---|---|---|---|---|---|---|---|---|---|
| | | | | | | | | | | | of Dmel\CG30413 (Hmel203003o: 4,365,922-4,366,814) |
| **BR** | | MSTRG.3484 | | 7.491 | -0.957 | 0.190 | Hmel203003o | 4363434 | 4364216 | Dmel\CG30413 | downstream 2488 bp of Dmel\CG30413 (Hmel203003o: 4,365,922-4,366,814), before MSTRG.3485 |
| **BR** | | MSTRG.3159 | HMEL036008g1 | 1.256 | 1.020 | 0.191 | Hmel203003o | 269089 | 271724 | Ubiquitin conjugating enzyme 84D | poor hit, E value 2.56 |
| **Chr 7** | | | | | | | | | | | |
| **RS** | **Day 5** | MSTRG.8012 | HMEL013683g1 | 0.642 | 3.345 | 0.162 | Hmel207001o | 8627421 | 8630143 | Gemin 3 | na |
| **RS** | | MSTRG.8188 | HMEL037634g1 | -0.737 | 1.789 | 0.162 | Hmel207001o | 9711504 | 9713128 | RNA polymerase III subunit G | na |
| **RS** | | MSTRG.8324 | HMEL012321g1 | 0.694 | 5.347 | 0.186 | Hmel207001o | 11485564 | 11535901 | ADP ribosylation factor-like 4 | na |
| **RS** | | MSTRG.8164 | HMEL013901g1 | 0.656 | 6.590 | 0.194 | Hmel207001o | 9458675 | 9505412 | Phosphoinositide-dependent kinase 1 | na |
| **RS** | **Day 7** | MSTRG.8186 | HMEL002406g2, HMEL037635g1 | 1.084 | 3.409 | 0.066 | Hmel207001o | 9708171 | 9719190 | Dmel\CG32681 | poor hit E Value = 9.44 |
| **RS** | | MSTRG.8049 | HMEL007780g1 | -1.426 | 1.081 | 0.144 | Hmel207001o | 8824630 | 8833804 | ringmaker | na |
| **RS** | | MSTRG.8321 | | -4.665 | -0.081 | 0.192 | Hmel207001o | 11474783 | 11476066 | inaF-D | poor hit , E value 9.86, in region of no genomic features, downstream 9293 bp of Hmel207001o: 11,485,359-11,485,712 |

| RS | | MSTRG.8258 | HMEL037660g1 | 0.480 | 3.766 | 0.196 | Hmel207001o | 10575157 | 10595319 | Dmel\CG8243 | na |
|---|---|---|---|---|---|---|---|---|---|---|---|

Table S17. Genes differentially expressed (FDR<0.05) between *H. melpomene cythera* wing regions within the QTL intervals.

| Phenotype | Stage | gene id | annotated gene | logFC | logCPM | FDR | scaffold | start | stop | Protein hit (Flybase) | NCBI blastp hit |
|---|---|---|---|---|---|---|---|---|---|---|---|
| **Chr 3** | | | | | | | | | | | |
| **BR** | **Day 7** | MSTRG.3151 | HMEL021694g1,HMEL021694g2 | -0.793 | 4.336 | 0.025 | Hmel203003o | 159359 | 167643 | Dmel\CG9701 | PREDICTED: lactase-phlorizin hydrolase-like [Papilio machaon] |
| **BR** | | MSTRG.3338 | HMEL021575g1 | -0.530 | 6.252 | 0.091 | Hmel203003o | 1912342 | 1917684 | tracheal-prostasin | trypsin-1-like [Vanessa tameamea] |
| **Chr 7** | | | | | | | | | | | |
| **RS** | **Day 5** | MSTRG.8196 | HMEL015765g1 | -1.024 | 5.557 | 0.024 | Hmel207001o | 9752688 | 9775897 | Leucine-rich repeat | F-actin-uncapping protein LRRC16A isoform X2 [Pararge aegeria] |
| **RS** | **Day 7** | MSTRG.8135 | HMEL021853g1 | 0.702 | 6.947 | 0.020 | Hmel207001o | 9318579 | 9319970 | Dmel\CG18294 (poor hit) | cuticle protein 18.6-like [Bicyclus anynana] |
| **RS** | | MSTRG.8147 | HMEL003671g1 | 0.438 | 8.687 | 0.174 | Hmel207001o | 9391192 | 9391719 | Dmel\CG13063 (poor hit) | cuticular protein hypothetical 9 precursor [Bombyx mori] |
