## Supplementary methods for "The genetic basis of structural colour variation in mimetic *Heliconius* butterflies"

**Supplementary Methods and results**
**The genetic basis of structural colour variation in mimetic *Heliconius* butterflies**

Melanie N. Brien, Juan Enciso Romero, Victoria J. Lloyd, Emma V. Curran, Andrew J. Parnell, Carlos Morochz, Patricio A. Salazar, Pasi Rastas, Thomas Zinn, Nicola J. Nadeau

### Supplementary Methods

#### *Sampling*

Non-iridescent *H. erato demophoon* and *H. melpomene rosina* were collected from Gamboa, Panama (9.12ºN, 79.67ºW). Iridescent *H. erato cyrbia* and *H. melpomene cythera* were collected from the area around Mashpi, Ecuador (0.17ºN, 78.87ºW). Within a few hours of emergence, wings of offspring were removed and stored in glassine envelopes and bodies were stored in NaCl saturated 20% dimethyl sulfoxide (DMSO) 0.25M EDTA solution to preserve the DNA. Many samples analysed in this study were previously used in Brien et al. (Brien *et al.*, 2018) and Bainbridge et al. (Bainbridge *et al.*, 2020).

#### *DNA extraction and sequencing*

Genomic DNA was extracted from approximately half of the thorax of each individual using the Qiagen DNeasy Blood & Tissue kit. Single-digest Restriction site-associated DNA (RAD) library preparation and sequencing were carried out by the Edinburgh Genomics facility. DNA was digested with the enzyme *PstI*, and libraries were sequenced on an Illumina HiSeq 2500 producing 125bp paired-end reads. Parents of the crosses (16 *H. erato* and 9 *H. melpomene*) were included at 2x higher concentration within the pooled library to produce a higher depth of coverage.

*Data processing and linkage map construction*

Sequence data processing and linkage map construction are described in detail in Bainbridge et al. (Bainbridge *et al.*, 2020). Briefly, RAD sequences were demultiplexed using the RADpools function in RADtools v1.2.4 (Baxter *et al.*, 2011). FASTQ files were mapped to the *H. erato* v1 reference genome (Van Belleghem *et al.*, 2017) or the *H. melpomene* v2.5 reference genome (Davey *et al.*, 2016) using bowtie2 v2.3.2 (Langmead and Salzberg, 2012). BAM files were then sorted and indexed with SAMtools v1.3.1 (Li, 2011) and PCR duplicates were removed using Picard tools MarkDuplicates v1.102. SAMtools mpileup and Lep-MAP data processing scripts (Rastas, 2017) were used to call genotype posteriors. Linkage maps were constructed with Lep-MAP3 (Rastas, 2017), using the modules ParentCall2 (with options ZLimit=2 and removeNonInformative=1) (Rastas, 2017), Filtering2, SeparateChromsomes2 (using sizeLimit = 50 and a LOD score of 12 for *H. erato* and 23 for *H. melpomene*) and JoinSingles2All. Finally, the module OrderMarkers2 was used to order the markers along the linkage groups, taking into account the genomic positions of the markers. Female recombination was set to 0 and male recombination set to 0.05 (following Morris et al. (Morris *et al.*, 2019)). We used LMPlot to visualise the maps and check for errors in marker order. Any erroneous markers that caused long gaps at the beginning or ends of the linkage groups were manually removed. Genotypes were phased using Lep-MAP's map2genotypes.awk script, and markers were named using the map.awk script and a list of the SNPs used to provide the scaffold name and position. The final linkage maps contained 5648 markers for *H. erato*, and 2163 markers for *H. melpomene*.

*Measurements of colour variation*

Wings were photographed under standard lighting conditions (full details in Brien et al. (Brien *et al.*, 2018)). A colour checker in each photograph was used to standardise the photographs using the levels tool in Adobe Photoshop (CS3). RGB values were extracted from two areas of each wing (proximal areas of both the forewing and hindwing) and averaged. Blue-red (BR) values were used as a measure of blue iridescent colour. These were calculated as (B-R)/(B+R), where +1 is completely blue and -1 is completely red. Luminance measured overall brightness and was calculated as R+G+B, with each colour having a maximum value of 255.

*Ultra small-angle X-ray scattering (USAXS)*

Ultra small-angle X-ray scattering (USAXS) data was generated at the ID02 beamline at the European Synchrotron (ESRF, Grenoble, France). The X-ray beam wavelength was λ = 1 Å and it had 12.45 keV energy. The beam was collimated to a section of 50 µm × 50 µm to probe a small area within a single scale. The scattered radiation was recorded with a high-sensitivity FReLon 16M Kodak CCD detector with an effective area of 2048 × 2048 pixels and a 24 µm pixel size, a sample to detector distance of 30.7m, accessible q range between 8.5x10-4 $nm^{-1}$ and 3.8x10-2 $nm^{-1}$, and limited interpretation of structure spacing to above 160 nm.

*Scanning electron microscopy (SEM)*

SEM on a small subset of individuals was used to confirm our scale structure measures from the USAXS data.  We sampled four iridescent *H. e. cyrbia* and three *H. m. cythera*, and eight *H. e. demophoon* and three *H. m. rosina* with black pigment colour. Representative SEM images of dorsal cover scales for each of the four races used are in Figure S8. First, we compared ridge spacing of blue individuals vs. black individuals on the dorsal side. This allowed us to gauge the expected range of scale structure variation in blue and black coloured scales. Secondly, we compared ridge spacing of dorsal vs. ventral scales within each race. This comparison allowed us to separate signals coming from overlapping scales in the USAXS experiment.

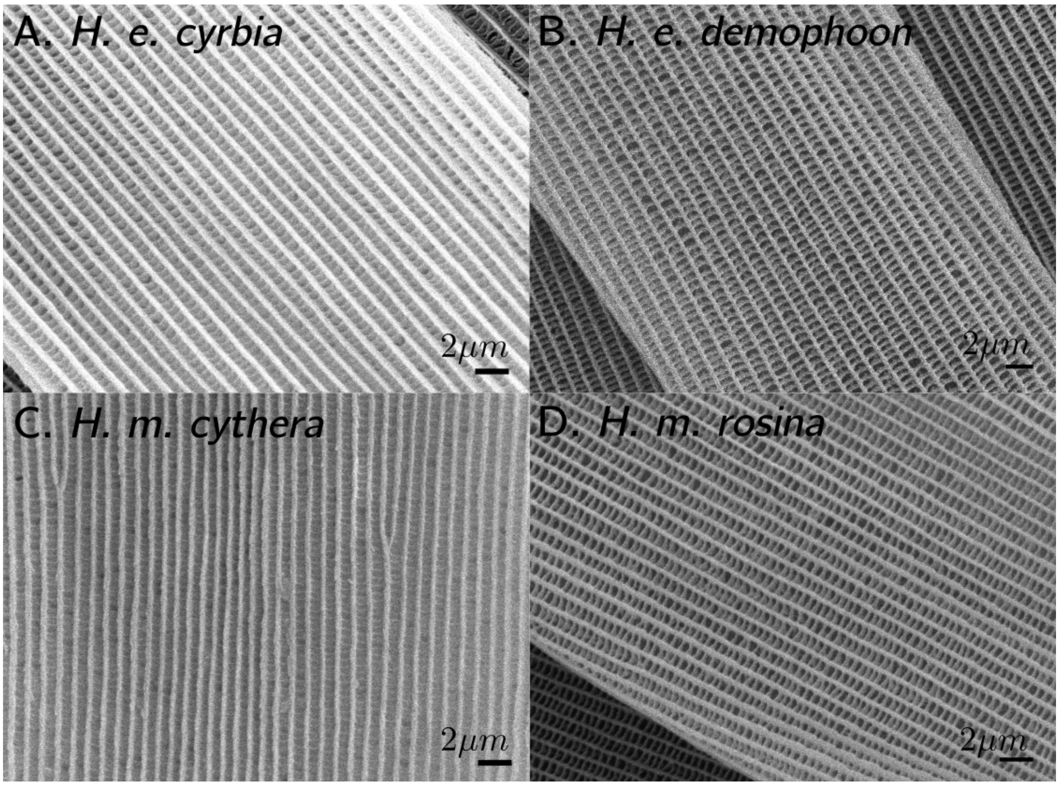


Figure S8: Representative SEM images of dorsal cover scales of iridescent blue (A) and matt black (B) races of *H. erato* and *H. melpomene* (C iridescent, D matt black).

Wing samples were cut from the region of interest shown in Figure S4. We mounted samples on aluminium SEM stubs using 9mm adhesive carbon tabs and coated them with a thin layer of gold using vacuum evaporation. Samples were imaged on a JEOL JSM-6010LA microscope and we used the InTouchScope software to take images of the regions of interest at a magnification level of 900X. Images included both cover and ground scales.

At the midpoint of the scale length, we drew a linear transect perpendicular to the orientation of the ridges (Figure S9). Ridge spacing was calculated using the PeakFinder tool (Vischer, 2013); a plugin for the Fiji image analysis package (Schindelin *et al.*, 2012), using a tolerance level of 60 and a minimum distance threshold of 10 pixels between peaks. PeakFinder detects spikes of intensity along the linear transect, each peak corresponding to a single ridge. It then yields the distances, in pixels, between subsequent peaks (Figure S9). Using the scale bar of SEM slides we determined the variable rate of conversion from pixels to nm to be $25 \times 10^{-3}$ / M where M is the level of magnification and we used this variable rate to convert distances between ridges to nm and average all distances between ridges to obtain average ridge spacing per scale.

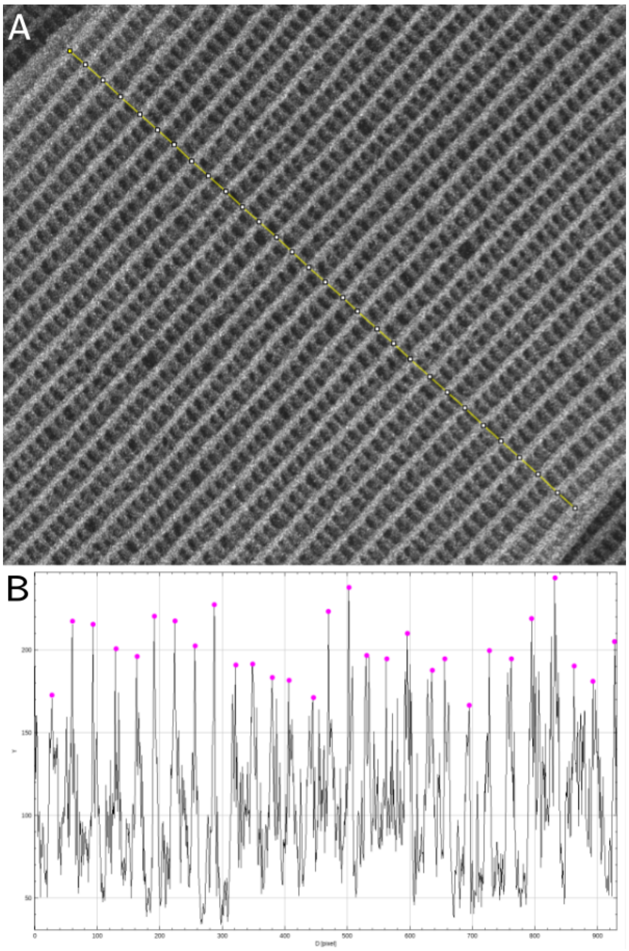


Figure S9: Measuring ridge spacing. (A) linear transect perpendicular to the orientation of the ridges. (B) peak finder output.

*Analysis of USAXS data to extract ridge spacing values*

We first did a qualitative joint assessment of our USAXS and SEM data to get an idea of the scale structures we captured in our experiment and how to best get their information. Our interpretation of scale structures in a USAXS scattering pattern is shown in Figure S10. Two important observations come from this joint assessment. Firstly, it can be observed that blobs of scattered radiation distribute along linear paths from the centre of the picture with equal spacing between them (Figure S10 A). This periodic scattering can be understood as the signature of the interaction of the beam with the array of parallel ridges on the surface of the scale (Figure S10 B) (Sivia, 2011). Secondly, the beam goes through the dorsal and ventral sides of the wing, projecting onto the detector, with the scattering due to structures of as many as four scales: cover and ground scales on both dorsal and ventral sides of the wing, in the case that all of them overlap. This can be observed in Figure S10 A, where at least three overlapping scales were picked up by the beam. Our SEM analysis revealed that scales with smaller ridge spacing are found on the dorsal side compared to the ventral side of the wing regardless of the structural colour of the butterfly analysed (Figure S11). These differences in ridge spacing between dorsal and ventral scales should be discernible in the USAXS data.

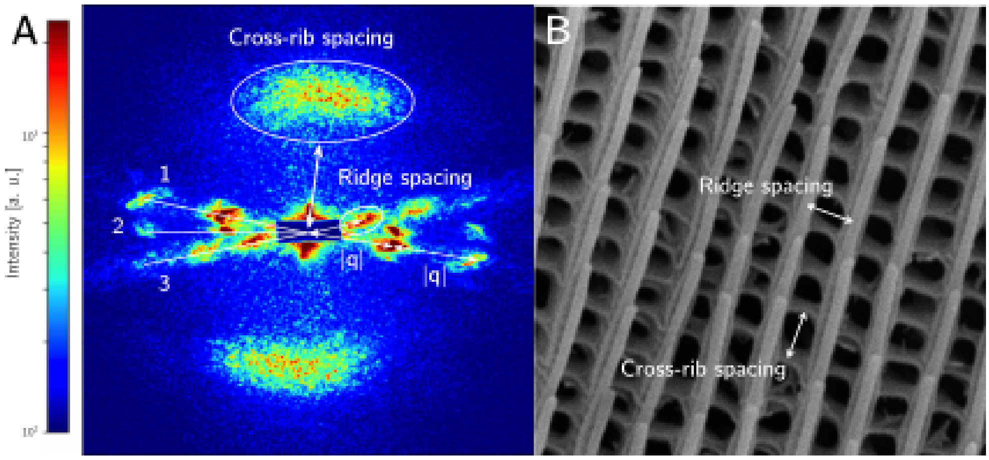

Figure S10: Representative scattering pattern captured in the USAXS experiment (A) and SEM slide of a *Heliconius* butterfly scale (B). Scattered intensities reveal oriented structures; these correspond to spacing between ridges and spacing between cross-ribs, which have a perpendicular orientation in the scale (B). For the ridges, it can be observed that blobs of scattered intensity appear with a regular spacing |q| along a linear path; this is due to their organisation as a parallel array in the upper lamina of the scale. Scattering of at least 3 different scales is observed, as indicated by the enumeration on the left hand side of the image.

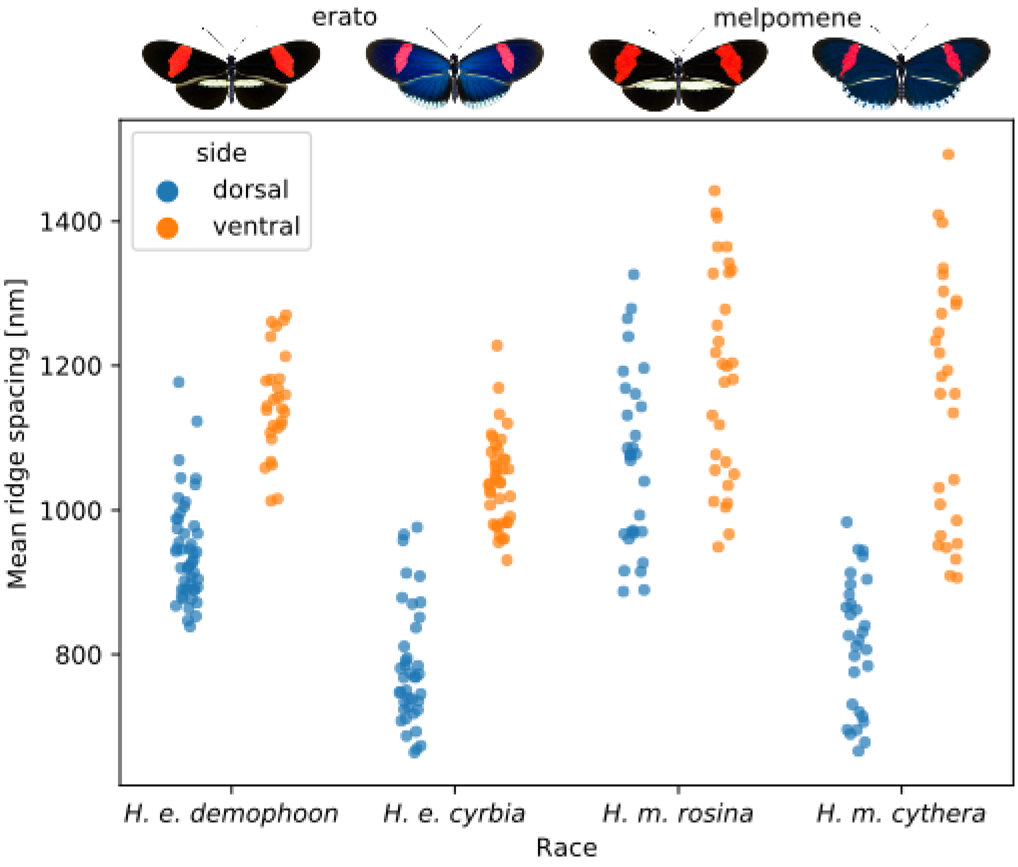


Figure S11: Comparison of ridge spacing between dorsal and ventral scales of insectary reared matt-black and iridescent blue *H. erato* and *H. melpomene* estimated from SEM images. Ventral scales show higher average spacing between ridges regardless of the presence or absence of iridescence in the wing.

We used these two observations to implement a script that extracts relevant scale structure information from the USAXS data. The script scans the 2D scattering pattern in non-overlapping regions of 5°. If the region contains periodic scattering it is assumed to correspond to the ridges of the scale and is kept for comparison with other sections that also show periodic scattering. The script keeps the section with the highest magnitude of the scattering vector *q* based on the observation that *q* should be larger for dorsal scales because they have tighter ridge spacing. The chosen region is recorded and the scanning section is rotated 85° anti-clockwise to capture the scattering due to cross-ribs, which are perpendicular to ridges in the scale. Azimuthal integration is then performed on the two regions of interest, reducing them to a one-dimensional representation of scattered intensity *I* as a function of the magnitude of the vector *q* in a similar way to previous studies (Brien *et al.*, 2018; Parnell *et al.*, 2018), applying corrections for transmitted flux and masking the pixels corresponding to the beam stop. The script uses the pyFAI (Kieffer and Karkoulis, 2013) and fabio (Knudsen *et al.*, 2013) libraries to do the azimuthal integration and USAXS data handling.

Despite dividing the scattering pattern in small regions to pick apart the signals due to different structures and scales, some frames still contained information of more than one scale. This happens in the case that overlapping scales have highly similar orientations (i.e. their ridges are parallel or nearly parallel). Thus, a visual inspection was done and we filtered out 1D scattering patterns that contained signals from more than one scale. We also aimed to reduce noise in the scattering signals to improve estimation of dimensions of scale structures. We ran a PCA on the 1D scattering data and retained PCs that cumulatively explained 90% or more of the variation in the data. The original data were projected back onto the retained PCs. The choice of threshold for the percentage variance explained is arbitrary, but we found 90% to be a sensible balance to retain signal but reduce noise.

The retained 1D scattering patterns were fitted to a Lorentzian curve (Figure S12) using the FitManager module of the silx package (Vincent *et al.*, 2020). The “center” parameter of the fitting procedure gives the magnitude of the scattering vector q. This magnitude contains information on the distance between peaks of intensity in reciprocal space. To get estimates of structure sizes in real space, we use the expression d = 2π/q where d is the distance between ridges or cross-ribs in real space. We averaged estimates of each structure on a per-individual basis to get estimates of average ridge spacing and average cross-rib spacing per butterfly.

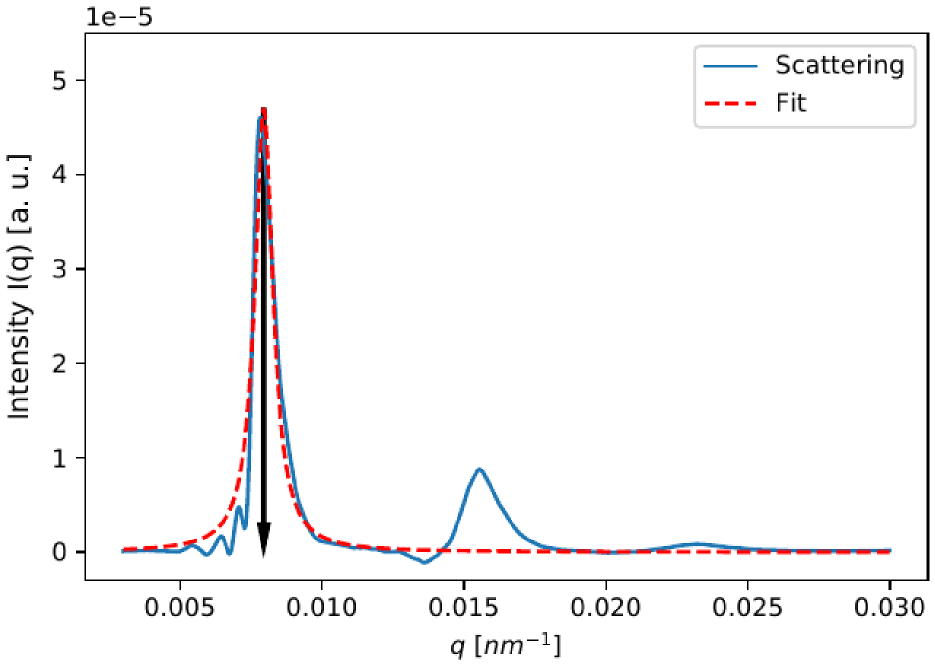


Figure S12: Example of peak fitting to find the magnitude of the scattering vector (q). 2D patterns are previously scanned for Bragg peaks and integrated over a small (5°) arc to produce the data of the blue curve. The fitting is done on the main peak using a Lorentzian curve (red). The peak intensity is observed at q ≅ 0.0075 (black arrow).

*Association analysis for H. melpomene*

The test was done in two steps: in the first step a mixed model is used to get estimates of trait values after adjusting for covariates and random polygenic effects using the linear model:

$$Y_i = \mu + \sum_j \beta_j C_{ji} + G_i + e_i \quad (1)$$

In this model we used a genomic kinship matrix to account for pedigree structure. In the model $Y_i$ is a vector of phenotypes for each individual, μ is the mean phenotype for the population, $C_{ji}$ is the value of covariate j for individual i, $\beta_j$ is the estimated effect for each covariate, $G_i$ are random additive polygenic effects and $e_i$ are the residual effects for each individual. The result of this first part are the trait residuals after accounting for covariates and random polygenic effects and is denoted by $\hat{e}_i$. The estimates of trait variation that were not explained by kinship or covariates in the first step are then fit to a second model:

$$\hat{e}_i = \mu + \beta g + e_i \quad (2)$$

In the second step, β is the parameter for the effect of each SNP tested, g is a single SNP and $e_i$ is the residual variation in phenotype for each individual i. We did the association tests on two sets: the first set included all *H. melpomene* families and two phenotypes; BR value and luminance. The second set included the EC70 family only and four phenotypes: BR value, luminance, ridge spacing and cross-rib spacing.

Genotypes were called using the 'mpileup' command in samtools v1.8 (Li, 2011) with minimum mapping quality set to 20, and the 'call' command in bcftools v1.8 (Li, 2011) run in consensus mode, with SNP p-value threshold set to 0.05 and calling only variant sites. For our association analyses we kept biallelic SNPs only. We applied the following quality control criteria to our SNP data: we excluded individuals and sites that were not at least 95% genotyped. Sites were also excluded from our analysis if they were not in Hardy-Weinberg equilibrium ($p<0.01$). The MAF threshold was set to the default value (5/2n, where n is the number of individuals). After quality control, we retained 146,122 markers genotyped across 211 individuals for the first set, and 63,132 sites genotyped across 69 individuals for the second set. We set sites called as heterozygous on the sex chromosome in females as missing (females are the heterogametic sex). Multiple testing was accounted for using two thresholds: A Bonferroni corrected threshold calculated as 0.05/k where k is the number of sites analysed, and a less conservative 5% false discovery rate threshold estimated using the GenABEL qvaluebh95 function (Aulchenko, de Koning and Haley, 2007).

*RNA sequencing data generation and processing*

RNA was extracted using QIAgen RNeasy. Paired-end sequenced data was generated from Illumina TruSeq RNA libraries (*H. melpomene* libraries were made using a TruSeq Stranded mRNA-seq kit and *H. erato* libraries were made using a TruSeq v2 kit) on 6 HiSeq v4 lanes. Low quality reads (PHRED <33) were removed and Illumina adapters clipped (using the ILLUMINACLIP option) using Trimmomatic v. 0.38 (Bolger, Lohse and Usadel, 2014). Read quality was then inspected using FastQC v. 0.11.8 (Andrews *et al.*, 2012) and the results aggregated and viewed using multiQC v. 1.5 (Ewels *et al.*, 2016). Trimmed reads were aligned to the respective *Heliconius* reference genomes using HISAT2 v. 2.1.0 (Kim *et al.*, 2019). HISAT2 was run with the '--known-splicesite-infile' option to enable alignment of reads with small anchors. Exon and splice site information was extracted from the respective species gene annotation files downloaded from Lepbase (v4) (Challis *et al.*, 2016). Gffread was used to convert the downloaded '.gff' file format into the required '.gtf format'. The python script 'hisat2_extract_splice_sites.py' was used to create the list of known splice sites to pass to HISAT2. In addition, HISAT2 was run with the -dta option to adapt the alignments for downstream programs, including StringTie. The *H. melpomene* reads were run with the –rna-strandedness RF option to account for the strandedness of the reads.

The output SAM files produced from HISAT2 were sorted and converted into BAM files using samtools v. 1.9 (Li, 2011). The average number of reads per sample was 18.2 million for *H. erato* and 15.3 million for *H. melpomene* (Figure S13 A). The average overall alignment to the respective species reference genomes was 76 % for *H. erato* and 78.0 % for *H. melpomene* (Figure S13 B).

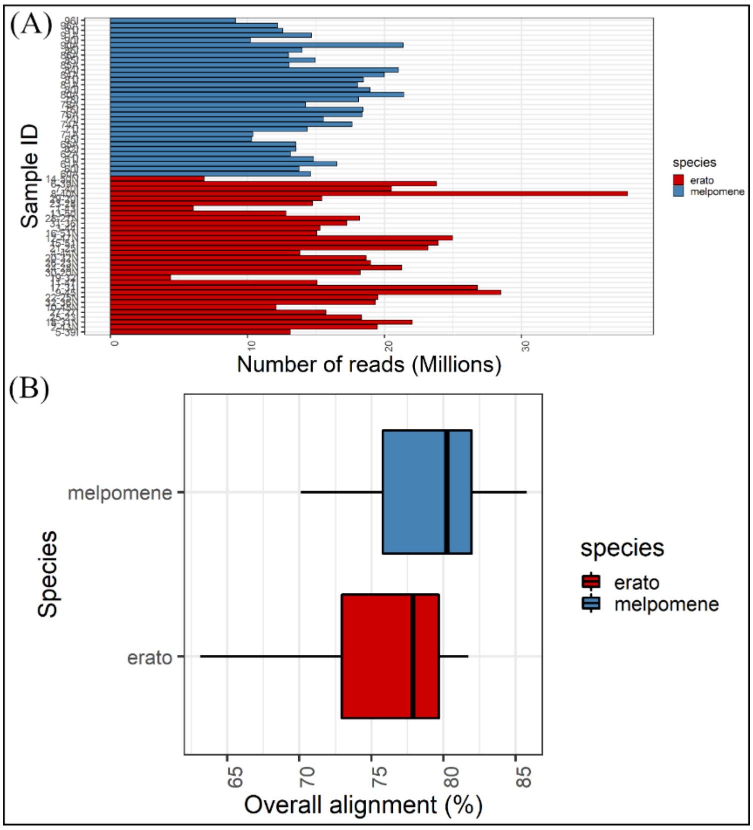


Figure S13: Filtering and alignment of RNA-seq reads of *H. erato* and *H. melpomene*. (A) Number of raw RNA reads in the samples of *H. erato* and *H. melpomene* in millions. (B) Overall mean percentage alignment of the RNA reads with the respective species genomes using HISAT2.

*Differential Expression analysis*

Differential expression analysis was performed using the R/Bioconductor package EdgeR v. 3.28.1 (Robinson, McCarthy and Smyth, 2010). Genes with low expression levels were filtered out from subsequent analyses using the 'filterByExpr' function (Chen, Lun and Smyth, 2016). Normalisation of library size was then performed using the trimmed mean of M-values (TMM) method (Robinson and Oshlack, 2010). Trended estimates of dispersion were calculated and negative binomial generalised linear models fitted using the 'glmQLFit' function.

#### *Functional annotation*

Differentially expressed genes between iridescent and non-iridescent races were annotated using the respective species InterproScan and Blastp databases downloaded from Lepbase (Challis *et al.*, 2016). For all other annotations, genes were manually compared to the non-redundant NCBI protein database using the Blastp algorithm, as well as the FlyBase BLAST protein database (Thurmond *et al.*, 2018).

#### *Gene Set Enrichment Analysis (GSEA)*

GO (gene ontology) enrichment analysis was performed in R v. 3.6.2 using the R/Bioconductor package topGO v. 2.38.1 (Alexa, Rahnenführer and Lengauer, 2006). Custom annotations were constructed using GO terms extracted from the InterproScan files downloaded from Lepbase. The gene universe comprised of all the genes expressed in the wings at 5 and 7 DPP taken from the respective species gene count matrix produced by StringTie. GO analyses were performed on the differentially expressed genes between the iridescent and non-iridescent which had an FDR $< 0.05$. A Fisher exact test was used to conduct the test using the 'weight01' algorithm. GO terms were considered significant when $P < 0.01$. GO enrichment circle plots were constructed using the package GOplot v. 1.0.2.

#### *Identification of convergent differentially expressed genes*

BLAST v. 2.9.0 (Altschul *et al.*, 1990) was used to identify orthologous genes between *H. erato* and *H. melpomene*. A blast database was created using reference protein sequences for *H. melpomene* downloaded from Lepbase (v4) (Challis *et al.*, 2016) using the 'makeblastdb' function. Then, using blastp protein sequences for *H. erato*, were aligned against the *H.*

*melpomene* reference to create a table of orthologous genes between the two species. For both 5 and 7 DPP, genes which were differentially expressed (FDR < 0.2) between iridescent and non-iridescent races in both *H. erato* and *melpomene* were identified. Genes were further filtered based on concordant expression patterns, i.e. only genes which were upregulated or downregulated in both *H. erato* and *H. melpomene* were kept. In addition, the analysis above was repeated using the DEG's between the iridescent and non-iridescent wing region of the iridescent species (FDR < 0.2).

### Supplementary Results

#### *Analysis of sexual dimorphism*

We tested for sexual dimorphism in the four measured traits (BR colour, luminance, ridge spacing and cross-rib spacing) and did not find any significant differences between male and female *H. melpomene* offspring from the largest brood (EC70) in all four traits (Table S18). Sexual dimorphism in *H. erato* was analysed in our previous study, finding differences in all four phenotypes in offspring with a maternal *cyrbia* grandfather (Brien *et al.*, 2018). Our sample size from the parental subspecies is small but did not reveal any large difference between the sexes for any of the traits measured, suggesting that the difference we see between the sexes in the crosses are largely due to sex-linkage, as we previously proposed (Brien *et al.*, 2018). Nevertheless, we included sex as a covariate in all our subsequent analyses to control for the possibility of slight sexual dimorphism.

Table S18: Comparisons between females and males for the four phenotypes in *H. melpomene*.

| | Females | Males | Welch's t | d.f. | p |
|---|---|---|---|---|---|
| Ridge spacing (nm) | 929.36 ± 6.44 | 929.72 ± 8.16 | -0.034 | 73.61 | 0.97 |
| Cross-rib spacing (nm) | 470.59 ± 3.16 | 468.29 ± 3.17 | 0.517 | 75.82 | 0.60 |
| BR | -0.21 ± 0.02 | -0.17 ± 0.02 | -0.988 | 75.25 | 0.32 |
| Luminance | 74.50 ± 2.84 | 70.01 ± 3.10 | 1.066 | 75.91 | 0.28 |

*Phenotypic variation*

Based on current understanding of the reflective structures in *Heliconius* and other butterflies (Wilts *et al.*, 2017; Parnell *et al.*, 2018), we predict that reduced ridge spacing should increase the brightness of the reflectance from the wing due to the increased density of reflective structures (ridges) on the wing. However, there are other aspects of scale nanostructure that we would also expect to affect colour hue and brightness, particularly lamellae layering. These aspects could correlate with ridge spacing if they are controlled by the same genetic and developmental mechanism, or they could be controlled independently. The extent to which the different aspects of colour and scale structure are correlated in our broods gives an initial approximation of the extent of genetic correlation between these traits, and therefore the extent to which we are likely to find that they are controlled by the same or different genetic loci. For the *H. erato* family for which we have USAXS measurements, we found significant correlations between scale structure and colour measurements: ridge spacing has a negative correlation of moderate strength with both BR value (Spearman's rho = -0.411, n = 56, p = 0.0013) and luminance (Spearman's rho = -0.46, n = 56, p = 0.0002, Figure S2), as expected. We also found significant correlations between ridge and cross-rib spacing (Spearman's rho = 0.342, n = 56, p = 0.008) and between BR value and luminance (Spearman's rho = 0.341, n = 56, p = 0.008). Cross-rib spacing is negatively correlated with BR value (Spearman's rho = -0.325, n = 56, p = 0.012) but not with luminance (Spearman's rho = -0.18, n = 56, p = 0.17).

On the other hand, the *H. melpomene* family only showed significant correlations for BR value and luminance (Spearman's rho = 0.295, n =73, p = 0.008) but no correlations were found for BR value and ridge spacing (Spearman's rho = 0.0012, n = 73, p = 0.991), luminance and ridge spacing (Spearman's rho = -0.117, n = 73, p =0.306, Figure S2), ridge spacing and cross-rib spacing (Spearman's rho = -0.088, n = 73, p = 0.441), BR value and cross-rib spacing (Spearman's rho = 0.05, n = 73, p = 0.622) or luminance and cross-rib spacing (Spearman's rho = 0.13, n = 73, p = 0.226). This separation of ridge spacing and colour variation may have arisen in *H. melpomene* as a result of a higher number of generations of mixing in the brood we used (EC70) in comparison to the *H. erato* brood. However, it would also be consistent with a greater genetic correlation between multiple aspects of scale structure in *H. erato* than in *H. melpomene*.

*Association analysis for H. melpomene*

Treating the EC70 cross as a backcross is not ideal as it ignores potentially interesting genetic variation that is segregating from the mother (which phenotypically and genetically appeared to be an F2 individual), and could lead to artefacts. Therefore, we also conducted a SNP association analysis including the full set of *H. melpomene* broods (F2 + EC70) and for the EC70 family alone, using a family-based score test for association (Chen and Abecasis, 2007) and controlling for family structure. The GWAS of the EC70 family did not reveal significant genetic associations for either of the scale structure traits (Figure S5). The SNPs with the lowest p-values were found on chromosomes 1, 7, 17 and Z for ridge spacing, and on chromosomes 13 and Z for cross-rib spacing. Similarly, we did not find significant associations between SNPs and BR values in EC70, although the lowest p-values were found on chromosome 3 and chromosome 10, which aligns with the QTL results. We did find SNPs significantly associated with variation in luminance on chromosome 3, with a single SNP scoring above both conservative genome-wide Bonferroni corrected thresholds. When analysing all of *H. melpomene* broods together accounting for pedigree structure, we found significant associations for both colour traits (BR value and luminance) on chromosome 3. Stronger support for association was found for luminance in comparison to BR value. Overall, the association analysis results supported the QTL results, but did not reveal additional loci.

*Gene set enrichment analysis (GSEA) of gene expression data*

GO term enrichment was performed using topGO (Alexa and Rahnenfuhrer, 2020) to identify biologically relevant processes underpinning the DE genes in our comparisons between subspecies of *H. erato* and *H. melpomene*. In *H. erato*, the majority of enriched GO terms occurred at 7 DPP, where out of the 1043 DE genes, 8 GO terms were significantly enriched across the three GO domains (4 Molecular Function, 2 Cellular Component and 2 Biological Process). Focusing on Molecular Function (Figure S14), the most enriched GO term was ‘acyl-CoA dehydrogenase activity’. Acyl-CoA dehydrogenase activity is involved in the first step of fatty acid β-oxidation; however, the specific functional role of this process is difficult to ascertain as fatty acids are required for numerous roles including: energy metabolism, precursors for eicosanoids and pheromones, synthesis of waxes and phospholipids (Arrese and Soulages, 2010). Manual Blastp search results indicate a number of the genes annotated to the term ‘Acyl-CoA dehydrogenase activity’ are candidate enzymes in the β-oxidation step of the Lepidopteran pheromone biosynthesis pathway (Ding and Löfstedt, 2015). Indeed, many of the additional enriched GO terms could relate to pheromone biosynthesis, including ‘oxidoreductase activity acting on paired donors’ and ‘fatty acid beta-oxidation’.

Interestingly, the GO term 'Voltage-gated cation channel activity' was also significantly enriched at 7 DPP in *H. erato*, with the majority of associated genes involved in calcium channel activity. At 5 DPP there was only 1 significant GO term (amide binding) out of the 907 differentially expressed genes between the races (Figure S14).

In *H. melpomene,* there were no significantly enriched terms at either 5 or 7 DPP. This lack of enrichment is likely due to the low number of differentially expressed genes (203 genes at 5 DPP and 78 genes at 7 DPP). Moreover, a significant proportion of the genes in both the gene universe and DE gene list lacked GO annotation, further minimising possible gene sets which may be significant. For example, from the 30,720 genes in the *H. melpomene* gene universe, only 25.1 % have a current GO annotation of some description.

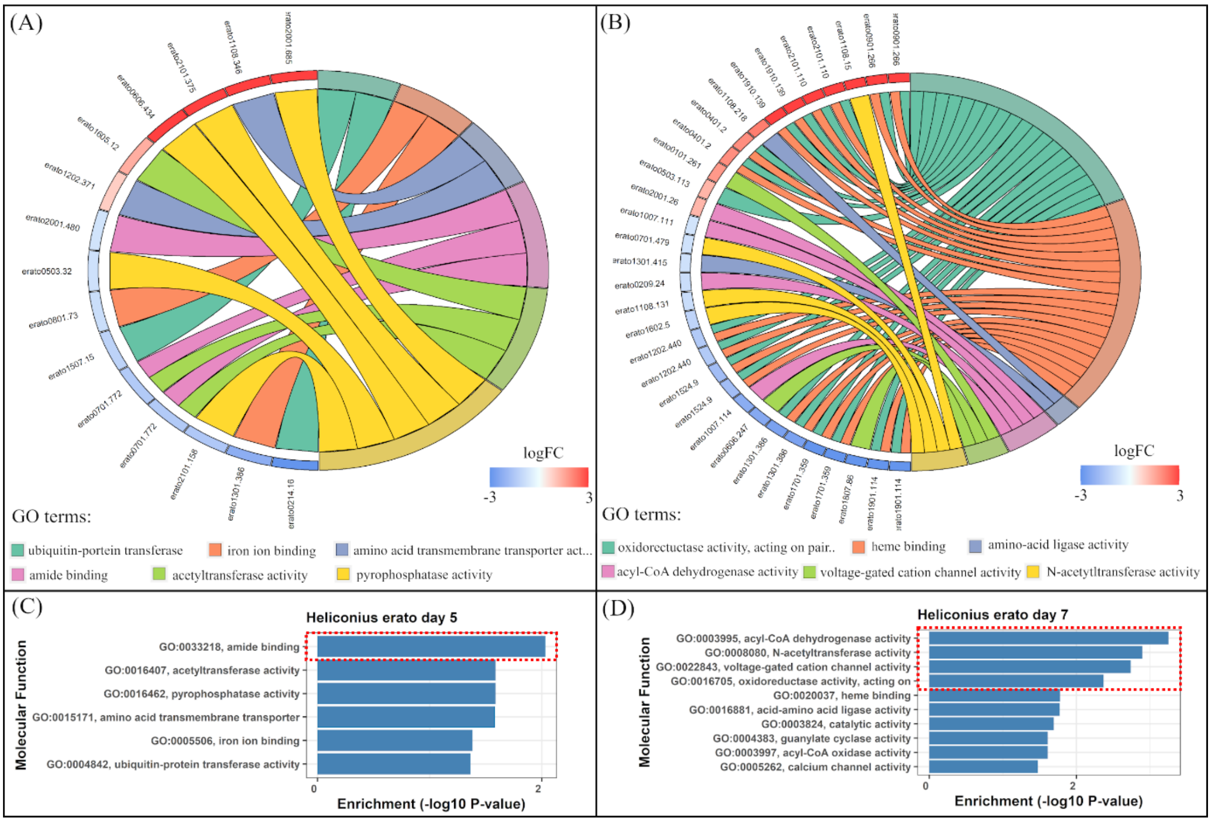


Figure S14: GSEA analysis (Molecular Function) of the differentially expressed genes between iridescent and non-iridescent races of *H. erato* at 5 and 7 DPP. (A-B) GO enrichment circle plots showing expression levels of genes belonging to the top 6 most enriched GO terms at 5 DPP (A) and 7 DPP (B). Genes associated with GO terms are on the left of the circle and associated GO terms are on the right. Colour scale of genes indicates amount of up/down regulation (LogFC). (C-D) Top enriched molecular function GO terms and descriptions for 5 DPP (C) and 7 DPP (D). All terms with an enrichment P-value greater than $-\log_{10}(2)$ are significant and are highlighted by a red dashed box.
